# Taking genomics outdoors: linking local adaptation, trait variation, and gene expression in grass ecotypes across a rainfall gradient

**DOI:** 10.64898/2026.01.04.697578

**Authors:** Jack Sytsma, Matthew Galliart, Kian Fogarty, Kori Howe, Bradley J. S. C. Olson, Sara G. Baer, David J. Gibson, Brian Maricle, Eli Hartung, Loretta Johnson

## Abstract

With increasing droughts, understanding local adaptation and drought tolerance in ecologically dominant species is crucial for enhancing ecosystem resilience. We leveraged a long-term reciprocal garden to assess local adaptation and drought responses in *Andropogon gerardi*, a foundation grassland species in the US Great Plains. Our objectives were to identify adaptive traits, explore gene expression responses across a rainfall gradient and under experimental drought and integrate trait-based analyses with gene expression profiles to test for local adaptation. Reciprocal gardens, established a decade ago, are composed of different *A. gerardi* ecotypes sourced along a rainfall gradient (MAP 480–1167mm yr^-1^) and sown as ecological communities. Rainout shelters imposed experimental drought. We hypothesized that ecotypes should perform best in their homesite, reflecting local adaptation. The dry ecotype should perform best under rainouts and exhibit traits and expression profiles favoring drought tolerance; the wet ecotype should favor traits and gene expression enhancing growth and resource acquisition. We found ecotypes had highest biomass and cover in their homesite, confirming local adaptation. Under experimental drought, the dry ecotype demonstrated improved performance at the wet site, confirming its adaptive value under water limitation. The dry ecotype showed stress-tolerance strategies (shorter, more water-efficient, upregulated drought-response genes), while the wet ecotype emphasized growth strategies (taller, higher biomass, upregulation of growth hormone gibberellin). Using co-expression networks, gene clusters linked adaptive traits, revealing genetic mechanisms of adaptation. Results advance our understanding of adaptation by linking gene expression and trait variation to drought responses in locally-adapted ecotypes, informing ecotype climate-matching under drought.

## 1. Introduction

Increasing frequency of drought has motivated research on how changes in rainfall affect the functioning of grasslands (Cook et al., 2015; Li et al., 2018; Griffin-Nolan et al., 2021). Drought is considered one of the most severe of environmental stresses to plants (Seleiman et al., 2021), therefore understanding plant tolerance to water limitation is particularly important (Smith et al., 2024). Furthermore, grasslands are sensitive to drought (Cherwin & Knapp, 2012; Knapp et al., 2015) and the impact of drought on grassland productivity has been underestimated (Smith et al., 2024). This sensitivity highlights the importance of understanding local adaptation as a mechanism of drought tolerance in grassland species, especially dominant grasses. Dominant grass species may play a crucial role in maintaining productivity (Zhang et al., 2025a) and ecosystem resilience in the face of changing rainfall (Van der Meer & Jansen, 2019).

Despite the well-documented importance of drought in grasslands (Cherwin & Knapp, 2012; Craine et al., 2013; Knapp et al., 2015), predictions of grassland functioning under drought are not well characterized in the context of the potential for species to adapt. This is despite the fact that intraspecific variation plays a key role in climate response by providing the raw material for adaptation (Westerband et al., 2021). Understanding local adaptation—where populations exhibit peak fitness in their home environments (Savolainen et al., 2007)—is essential to predict how plants respond to environmental variation and climate stressors (Savolainen et al., 2013).

Significant progress has been made by separately studying local adaptation in ecology (Leimu & Fischer, 2008; Wadgymar et al., 2022) and genomics (Savolainen et al., 2007; Kang et al., 2023) Yet, a critical gap remains in bringing genomics into ecological field experiments (Ungerer et al. 2008, Johnson et al. 2022, Sytsma et al., accepted). Bridging this gap is essential for understanding how genetic variation drives responses to environmental stressors under realistic, complex field conditions. By combining trait-based analyses with modern genomic approaches, we can both characterize phenotypic variation (Fortunato & Pires, 2020) and uncover gene expression profiles associated with ecotypic drought response (Pardo et al., 2020).

Several studies have identified drought-responsive genes in plants, especially in the model plant *Arabidopsis thaliana* (Lovell et al., 2015; Kang et al., 2023). However, knowledge of gene expression in wild plants is limited, particularly in long-lived perennial C_4_ grasses that endure long periods of drought (Weaver, 1968). Research on some grasses such as *Panicum* spp. (Lovell et al., 2016; Heckman et al., 2025) and *Brachypodium* spp. (Song et al., 2019; Decena et al., 2021) shows that substantial intraspecific variation in drought-responsive genes exists within species. Uncovering such intraspecific variation in stress response under field conditions is crucial for identifying traits that enhance drought resilience and inform restoration strategies to guide selection of climate-resilient seed sources (Torok et al., 2024). Many ecological genomic studies are conducted using single-species trials and most are short-term (Johnson et al. 2022, Sytsma et al., accepted). By contrast, our research platform, set in ecological communities, extends for over a decade, thus encompassing interannual rainfall variation.

We focus on local adaptation in big bluestem (*Andropogon gerardi* Vitman), a dominant C_4_ perennial grass that contributes up to 70% of aboveground biomass in tallgrass prairies (Knapp et al., 1998) and can disproportionately shape community and ecosystem structure (Mendola et al., 2015; Ren et al., 2023). This species spans a steep rainfall gradient across the U.S. Great Plains (mean annual precipitation MAP 480–1167 mm yr^-1^) putatively resulting in dry, mesic, and wet ecotypes (Gray et al., 2014; Johnson et al., 2015; Galliart et al., 2019, 2020). While *A. gerardi*’s responses to rainfall are well studied (Fay et al., 2003; Slette et al., 2023), its drought tolerance at the transcriptomic level remains poorly understood. To address this, we used an existing reciprocal garden experiment, established in 2009 across the rainfall gradient, with ecotypes cross-transplanted planted in prairies and part of the plots subjected to ∼50% rainfall reduction via rainout shelters (Johnson et al., 2015, 2022). This design enables testing for local adaptation under realistic ecological conditions. Measurements taken ten years post-establishment reflect long-term successional dynamics critical for evaluating adaptation in long-lived grassland perennials (Ren et al., 2023, 2024; Gibson et al., 2024).

The overall goal of this study was to assess local adaptation in *A. gerardi* ecotypes combining both trait-based and genomic analyses. To achieve this, the first objective was to evaluate phenotypic evidence for local adaptation using trait-based analyses to test whether phenotypic differentiation among ecotypes corresponds to environmental gradients and aligns with adaptive strategies under drought. We hypothesized that local ecotypes would perform best in their home environments, with reduced performance when transplanted to non-native sites, consistent with local adaptation. We expected the dry ecotype to exhibit a drought-tolerant strategy such as shorter stature and higher water-use efficiency (Grime, 1988; Jardine et al., 2021), while the wet ecotype would show traits related to growth and resource acquisition, including greater height and higher biomass (Grime, 1988; Moles et al., 2009; Jardine et al., 2021). Experimental drought was predicted to reduce biomass, photosynthetic rates, and delay flowering across all ecotypes, with the dry ecotype being least impacted.

Our second objective was to investigate transcriptomic signatures of local adaptation and their alignment with ecotypic strategies across the natural rainfall gradient and under experimental drought. Specifically, we aimed to test how gene expression differs and whether gene expression patterns reflect the functional adaptive strategies of each ecotype. We hypothesized that the dry ecotype would upregulate drought-response genes (e.g., DREB; Zhang et al., 2025b), while the wet ecotype would upregulate genes related to growth (e.g., gibberellin response; Shah et al., 2023) and resource acquisition. Additionally, we expected that rainout treatments would induce stress-related genes, including heat shock proteins (Rahman et al., 2022) and ABA-regulated genes involved in stomatal control (Aimar et al., 2014).

Our final objective was to uncover genomic basis of local adaptation by integrating both phenotypic and transcriptomic data. We used co-expression network analysis (Aoki et al., 2007; Serin et al., 2016) and hypothesized that specific gene modules would correlate with physiological traits. For example, we predicted that genes involved in ABA signaling would be co-expressed with stomatal conductance. Similarly, we expected that genes associated with growth, such as the growth hormone gibberellin (Shah et al., 2023), would be co-expressed with plant height.

This study is among the first to link local adaptation, trait variation, drought response, and gene expression profiles in realistic and long-term ecological communities across a broad rainfall gradient, offering a rare window into the role of local adaptation to rainfall. By combining trait-based analyses with genomic characterization, we identify genetic and physiological adaptations that enable dry-adapted ecotypes to thrive under water limitation. Our findings will offer critical guidance for matching seed sources to future climate conditions (Doherty et al., 2017), informing restorations that prioritize climate-adapted ecotypes to improve long-term ecosystem function under increasing drought frequency and intensity (Baer et al., 2019).

## 2. Materials and Methods

### 2.1. Focal Species

*Andropogon gerardi* is distributed across most of eastern North America (USDA Plants Database, 2022) spanning broad rainfall gradients and dominates in the center of the range (∼450-1250mm yr^-1^). In addition to its dominance as part of the tallgrass prairie, it is also an important forage for cattle, accounting for $8 billion USD of the US rangeland’s economy (USDA NASS, 2023). Furthermore, *A. gerardi* is widely used in grassland plantings through the USDA’s Conservation Reserve Program, which has converted over 3.2 million ha of cropland to grassland (CRP, 2023).

We used the newly annotated genome of *A. gerardi* (*Andropogon gerardi* Hap2 v1.1; Joint Genomics Institute, 2023) that enables gene expression analysis. With a 2.5B bp scaffold and 88,437 protein-coding genes, it offers a valuable resource for studying genomic molecular mechanisms of drought adaptation, crucial for restoration efforts and its role as forage in future droughts. Being evolutionarily closely related to the well-studied crop plants *Zea mays* and *Sorghum bicolor* (Swaminathan et al., 2010), this relationship allows us to take advantage of knowledge of their genomes and annotations.

### 2.2. Reciprocal Garden Establishment Along the Natural Rainfall Gradient

We utilized a long-standing reciprocal garden platform of seeded ecological communities established over a decade ago. Full details on plot establishment can be found in Johnson et al. (2015; 2022). Briefly, in 2008, seeds of *A. gerardi* were collected from four populations (native prairies) in each of three distinct regions along a precipitation gradient (Figure 1a; Table S1), which at the time defined regional putative climate ecotypes. Reciprocal transplant gardens were established in 2009 with each ecotype sown into plots in Colby, KS, Hays, KS, Manhattan, KS, and Carbondale, IL, USA (driest to wet sites, MAP 480-1167mm yr^-1^, Figure 1a). All plots were also sown with the same mix of nine native subordinate species representing multiple functional groups to assemble a community and simulate a restoration (Table S1; Johnson et al., 2015).

**Figure 1.**
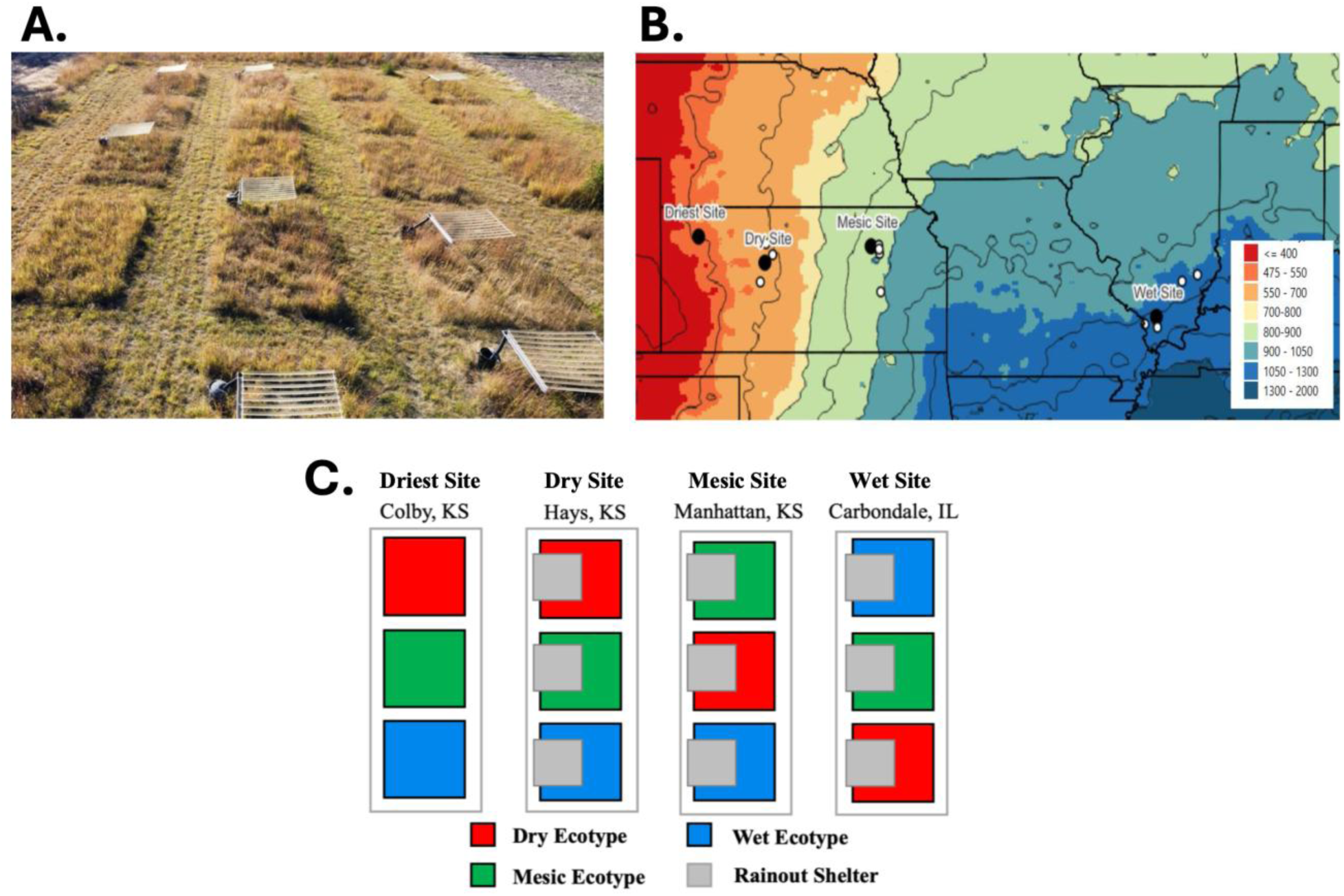
A. Aerial photograph of the mesic garden site, showing the spatial arrangement of plots within the prairies. This image highlights the ecological context of the experiment and illustrates how the reciprocal gardens are established within a grassland community. B. The location of the source populations, the garden sites along the natural rainfall gradient of the US Midwest. The location of reciprocal gardens plantings are black filled points and source populations are hollow white circles. Color gradients denote mean annual total precipitation in mm from 400-1250mm yr^-1^. C. Transplant design for garden plots with rainouts in gray and where ecotype plots (dry, mesic, or wet ecotypes) are indicated by colors (red, green, and blue). At each planting site, there are four replicate plots and plots were randomized at each site.

Colby KS had no local ecotype and was included to test ecotypic threshold to drier conditions as might be experienced in the future. Ecotypes were assigned to plots in each site according to a completely randomized block design where each block consisted of three 4 m × 4 m plots each sown with one of three ecotypes (Figure 1b). Plots were not weeded and volunteers from the regional species pool also established in the plots. The community aspects of the reciprocal garden platform are described elsewhere (Ren et al., 2023; 2024; Gibson et al., 2024).

To test for the specific role of rainfall in local adaptation, experimental rainouts on all ecotypes were established in 2011 in three of the sites (Hays, KS, USA, dry site-MAP 546 mm yr^-1^, mesic site-Manhattan, KS, USA, MAP 891 mm yr^-1^, and wet site-Carbondale, IL, USA, MAP 1256 mm yr^-1^, Figure 1b). Rainouts were not established in Colby KS, the driest site, because precipitation is already low there. Rainouts were designed to reduce ∼50% of ambient rainfall over half of each plot by using the design of Yahdjian and Sala (2002) and 2 m × 2 m roofs were installed in mid-June and taken down after the growing season each year. Rainfall runs into gutters and is then collected in rain barrels and disposed of away from the plots. At the wet site, using tubing connected to the gutter, excess water is drained away from the plots.

To assess soil moisture, we used METER volumetric soil moisture probes at 20cm depth (ECH20 Onset S-SMD-M005 Large Volume Soil Moisture Sensor; Meter Group, Pullman, WA, USA). Probes were installed in one block per site (n=4 per site) all three sites, including under rainouts. For Colby that did not have rainouts, we measured moisture in each block (total 28 probes). We took readings approximately once a week per block in ambient and rainout conditions at each site. Rainouts reduced soil moisture by ∼10-15% compared to ambient (Figure S1).

### 2.3. Objective 1: Testing for Local Adaptation Response Variables: Cover and Biomass

To assess local adaptation, we measured vegetative cover and aboveground biomass in 2021 and 2022 during peak season (late July to early August), over a decade after the establishment of the plots. Canopy cover served as a proxy for abundance, predicting success in long-lived perennials (Damgaard, 2015). For cover, we used 1m² quadrats with 81 intersections per plot and recorded *A. gerardi*, other grasses, forbs, or bare ground at each intersection. We also harvested aboveground biomass by clipping from 50 cm × 20 cm quadrats at the end of each growing season. For both cover and biomass, we used four non-overlapping quadrats per plot, for each of the three ecotypes in the four blocks at all four sites, including cover under rainouts (336 quadrats annually for cover and biomass).

#### Response Variables: Trait Measurements on Individual Plants

Early in the 2022 growing season, five individual *A. gerardi* plants were randomly selected from each plot and rainout treatment at each site and marked with flags. The driest site (Colby, KS, USA) did not have rainouts, so five plants were selected per each plot. We measured morphological, physiological, and reproductive traits on 420 individual plants in early, peak, and late (June, July, August) growing seasons. For brevity, here we include peak season measurements with early and late season results included as supplemental data.

##### Physiological Traits

To evaluate leaf physiological traits and gas exchange, we employed several techniques. We measured leaf chlorophyll absorbance with a SPAD-502 chlorophyll meter (Konica Minolta, Japan) of an average of at least three mature leaves. We used a LI6400-XT IRGA system (Li-Cor Biosciences, Inc., Lincoln, NE, USA) to assess gas exchange on sunny days between 10:00-14:00h. We measured photosynthesis, stomatal conductance, transpiration, water use efficiency (WUE, photosynthetic rate/ transpiration rate), and internal carbon dioxide. Gas exchange was measured on a fully expanded leaf with CO_2_ levels at 400ppm, humidity and temperature at ambient levels, and photosynthetically active radiation at 1500 μmol photons m^-2^s^-1^. Measurements were made when rates had stabilized with a coefficient of variation <1 for one minute.

##### Morphological Traits

To assess key plant traits and growth metrics, we employed several methods for measuring plant morphology and biomass. First, we determined height by extending leaves vertically from the ground to the tip of photosynthetic tissue. We measured plant diameter as an average of two orthogonal measurements of entire plant width. We assessed leaf width as an average of three mature leaves at their broadest section. We measured thickness of three mature leaves using a hand caliper (Mitutoyo Corp. Kawasaki Japan), excluding the midvein, and averaged these values per plant. In peak season (Julian days 177-192), we collected vegetative tissue from each plant at the midsection of 3-5 mature leaves to determine %C and %N. We harvested aboveground biomass in late fall by cutting individual plants at soil level, drying at 60°C, and weighing.

##### Reproductive Phenology

To track the phenological development of each plant, we assessed the date of bolting, defined as a clear presence of a flowering stalk, and the date of flowering, defined as the first presence of anthers. Plants were monitored throughout the season every 1-2 weeks until all flowered or until the date of first frost at each site.

### 2.4. Statistical Analysis: Cover, Biomass, and Functional Traits

To assess plot-level cover, biomass, and functional traits, we used a standard least squares regression model with a split-plot design in JMP Pro17 statistical software (SAS Institute Inc., Cary, NC, 2023). The main plot factors were site and ecotype, while the subplot factor was treatment (ambient or rainout), with all two-way and three-way interactions included in the model. For both cover/biomass and traits, site, ecotype, and rainout treatment (ambient or rainout) were fixed factors, and block number and quadrat/plant repetitions were treated as random effects. Separate models were constructed for early, peak, and late season data, as well as a repeated measures model to assess changes over time. When normality assumptions were violated, response variables were transformed using log transformations to improve the distribution of residuals. ANOVA and Tukey’s HSD tests were used to assess significance (p<0.05).

#### Ordination Analysis

We used a principal components analysis (PCA) to explore correlations of traits across ecotypes, sites, and rainout treatment. For different principal components analysis (PCA) were conducted separately under ambient conditions by ecotype, site, homesite, or treatment (ambient or rainout). All trait data were log-transformed to normalize distributions before they were used in correlation analysis and used a 95% confidence interval to show statistical significance.

### 2.5. Objective 2: Differential Gene Expression

#### Sample Collection and Extraction

Due to the strong phenotypic differences between ecotype and site extremes (dry and wet ecotypes, the driest, dry, and wet sites), we used these for gene expression analysis. We also analyzed gene expression profiles in ambient and rainout treatments. We collected 2-3 mature leaves from one plant of wet and dry ecotypes per block (n=40) at the same plants for which phenotypes were collected on the same day as traits from peak-season, allowing linkage between traits and gene expression. The leaves were flash frozen on dry ice in the field and transferred to -80°C freezer at the laboratory. Detailed methods on RNA extraction, sequencing, and analysis are found in Methods S1.

#### Differential Gene Expression and Identification of GO Terms

To identify differentially expressed genes (DEGs), we used DESeq2v3.112 (Love et al., 2014) on all pairwise comparisons using an adjusted probability (padj)<0.05. We categorized DEGs according to pattern of expression and up- or down-regulation over three comparisons (ecotype, site, rainout treatment). We identified genes using BLASTx (Camacho et al., 2009) to the *Zea mays* ensemble 58 and *Sorghum bicolor* rNCBlv3 reference genomes on the NCBI database at an *E*-value cut-off of 1 × 10^−6^ and selecting the top 20 hits.

Differentially expressed genes (|logFC| > 1, FDR-adjusted p-value) were visualized in a volcano plot (-log10 p-value vs. log fold-change) using the VolcanoPlot tool in Galaxy (https://usegalaxy.org/). The plots include only genes meeting the significance threshold (padj < 0.05; |logFC| > 1), highlighting those that are significantly upregulated or downregulated under different conditions. We visualized significant DEGs using a heatmap and clustered using “pheatmap” (Kolde et al., 2019) using R v1.0.12. Genes were hierarchically clustered by expression pattern using Pearson’s correlation, and red indicates upregulation while blue indicates downregulation.

We implemented Blast2GO 5.25 (Conesa & Gotz, 2008) to determine gene ontologies (GO terms), GO term enrichment, and annotations using 5% FDR cutoff. The complete lists were further filtered to focus on the GO terms that were within the top 10% of most significantly enriched within the original 5% FDR.

### 2.6. Objective 3: Linking Phenotype to Gene Expression

#### Weighted Gene Co-expression Network Analysis (WGCNA)

We used the WGCNA R package (Langfelder & Horvath, 2008) to identify significantly co-regulated gene cluster modules with phenotypic traits. Following module identification steps, we explored the correlation between module expression, ecotype (wet or dry), site (driest, dry, or wet), treatment (ambient or rainout), all two-way and three-way interactions, to the phenotype of the same plant from which the tissue was taken. Heatmaps were based on module eigengene values and included both correlation coefficients and corresponding p-values to assess the robustness of each association. Additionally, we used the R package “igraph” (Csardi & Nepusz, 2006) to construct network visualizations, illustrating the interactions between key genes and functional traits. Networks were built from adjacency matrices derived from module membership values (kME), and edges were filtered based on a minimum module membership threshold |kME| > 0.7, and significance of gene-trait correlations (padj < 0.05), retaining only the strongest connections. This filtering resulted in simplified network visualizations that highlight key hub genes and their putative functional associations with phenotypic traits.

## 3. Results

### 3.1. Objective 1: Ecotypic Variation and Local Adaptation

#### 3.1.1. Plot-Level Cover and Biomass

At the plot level in 2022, we found significant main effects of site, ecotype, and their interaction on *A. gerardi* canopy cover and biomass (all p<0.05), as well as a significant interaction of site × ecotype × treatment (p<0.05; Table S2). For 2021, patterns of biomass and cover mirrored the 2022 trends (Table S2b, Figures 2, S2).

**Figure.2.**
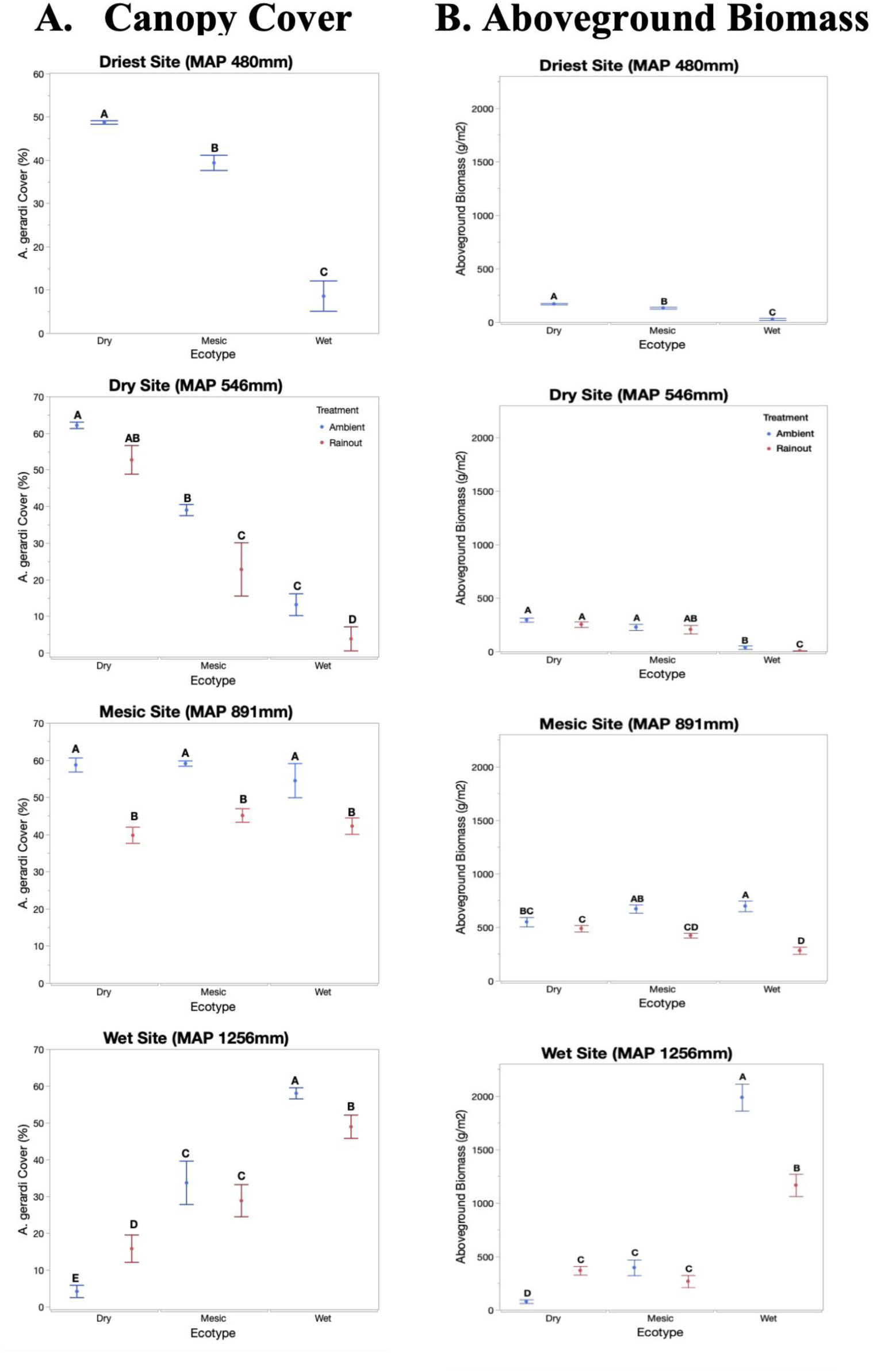
Comparisons of community-level response variables. A. canopy cover and B. aboveground biomass of *A. gerardi* in 2022. Ambient conditions are blue, and rainout is in red. The points indicate the mean, error bars represent standard errors, and the different letters denote significantly differences between ecotypes and treatments within a site and year (*α* = 0.05). The first year (2021) is presented in Figure S2.

##### Ecotype Variation by Site

We consider ecotype response at each site separately and focus mainly on 2022. Starting with the driest site (Colby, KS, USA), the dry ecotype had significantly higher cover (48.9% ±0.6) compared to the wet (8.8% ±1.2) and mesic (39.1% ±2.1) ecotypes (all p<0.05; Figure 2a). Moving on to the dry site (Hays, KS, USA), again the dry ecotype similarly had significantly higher cover (63.2% ±1.7) than the wet ecotype (15.4% ±1.8; p<0.05), and the mesic ecotype had intermediate values (Figure 2a). At the driest and dry sites, biomass showed similar trends in 2022 (Figure 2b). Progressing eastward to slightly wetter conditions, at the mesic site (Manhattan KS, USA), cover among ecotypes did not differ and were ∼60% (Figure 2a). Moving to the wet site (Carbondale IL, USA) the wet ecotype demonstrated significantly highest (60.6% ±1.5) cover, while the dry ecotype had only 6.2% ±2.0 cover, and the mesic ecotype was intermediate (34.7% ±4.6; Figure 2a). The mesic ecotype showed intermediate cover (40-60%) at all sites (Figure 2a). Biomass showed similar trends as cover in 2022 (Figure 2b).

For 2021, biomass and cover patterns mirrored the 2022 differences across sites (Figure S2).These patterns support evidence for local adaptation of the wet and dry ecotypes, with the dry ecotype demonstrating significantly highest cover and biomass at its dry home site and the wet ecotype showing significantly highest cover and biomass its wet home site (Figure 2).

##### Rainout

Regarding rainout treatment, at the dry site, mesic and wet ecotypes demonstrated significantly reduced cover under rainouts compared to the ambient (mesic ecotype ambient 40.4% ±0.7 vs rainout 26.2% ±3.2; wet ecotype ambient 15.4% ±1.8; vs rainout 4.8% ±0.9; Figure 2a). At the mesic site all ecotypes showed significantly reduced cover under rainouts (Figure 2a). At the wet site, the wet ecotype demonstrated a significant reduction in cover under rainouts compared to the ambient (60.6% ±1.5 vs 52.3% ±1.9; Figure 2a). Interestingly, also at the wet site, the dry ecotype had significantly higher cover under rainouts compared to ambient (18.7% ±3.1 rainout vs 6.2% ±2.0 ambient; Figure 2a). Results for biomass (Figure 2b) mirrored these trends. In 2021 under rainouts, biomass and cover generally showed similar trends as 2022 (Figure S2).

#### 3.1.2. Functional Trait Measurements

In terms of individual plant functional traits, at peak season, all 14 traits showed significant effects of ecotype and site; four out of 14 traits demonstrated significant main effects of rainout treatment (Table S2). The interaction between site x ecotype was significant (p<0.05) for all traits in all seasons, consistent with expectations for local adaptation. All other interactions and seasons are presented in Table S2.

##### Ecotypic Trait Variation by Sites

Across sites, wet and dry ecotypes exhibited distinct functional strategies, but these differences varied by site, often reflecting local adaptation. Since the mesic ecotype and mesic site showed intermediate trait values (Figures 3, S3), offering limited insight into divergent adaptive strategies, we focus our comparisons on the dry and wet ecotypes at the driest, dry, and wet sites.

**Figure 3.**
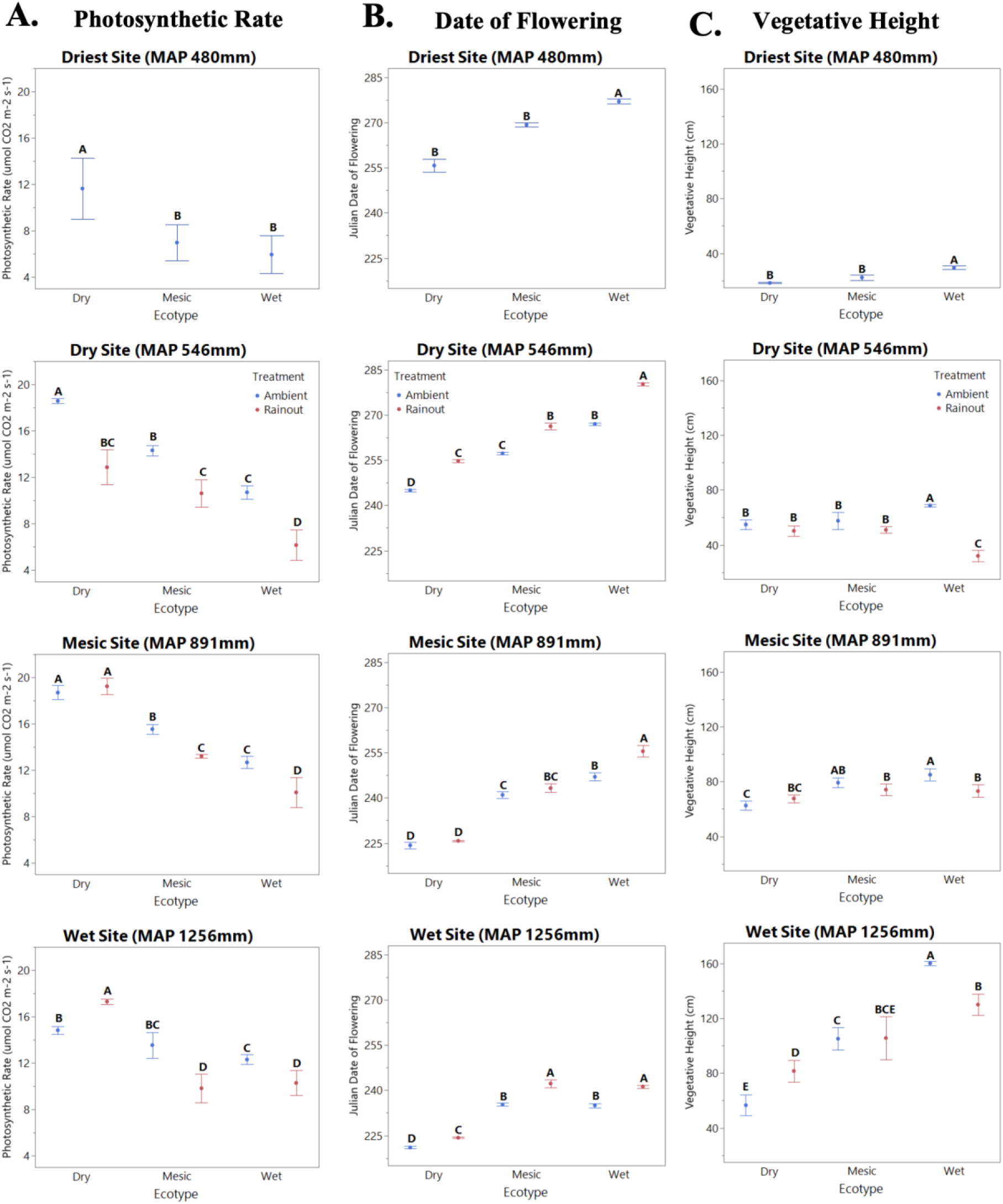
Comparisons of individual plant traits. A. photosynthetic rate, B. date of flowering, and C. vegetative height across sites (driest, dry, mesic, and wet) in peak season 2022. Ambient is blue, and rainout treatment is in red. The error bars represent standard errors, and the different letters denote significantly different groups among the comparisons within site (*α* = 0.05).

##### Driest Site

First, at the driest site, the dry ecotype compared to the wet ecotype demonstrated significantly greater photosynthetic rates (dry 11.2 µmol CO_2_m^-2^s^-1^±1.1 vs wet 5.0±0.2), higher stomatal conductance (dry 0.139 mol H_2_Om^-2^s^-1^±0.003 vs wet 0.09±0.01), greater water use efficiency (WUE; dry 3.64 µmol CO_2_/H_2_Om^-2^s^-1^±0.52 vs wet 1.05±0.04), higher nitrogen concentration (dry 1.54%±0.06 vs wet 1.31%±0.05), thicker leaves (dry 0.21 mm±0.02 vs wet 0.17±0.01) but lower transpiration (dry 2.78 µmol H_2_Om^-2^s^-1^±0.06 vs wet 4.28±0.06; all p<0.05; Figures 3a, S3). The dry ecotype bolted earlier than the wet ecotype by nine days (Julian date, JD dry 246.1±0.5 vs wet 255.3±2.9) and flowered earlier by 20 days (dry JD= 258.0±2.7 vs wet 278.3±3.0; Figures 3b, S3). However, the wet ecotype compared to the dry ecotype was nearly twice as tall (wet 32.4 cm ±5.6 vs 18.4±0.4), had wider leaves (0.74 cm±0.09 vs 0.60±0.01), and greater plant diameter by ∼30 cm (wet 53.8 cm±9.5 vs dry 21.4±1.6; all p<0.05; Figures 3c, S3).

##### Dry Site

Next, moving on to the dry site, the dry ecotype showed generally similar patterns as the driest site. The dry ecotype, at its homesite, compared to the wet ecotype showed significantly greater photosynthetic rates (dry 19.4 µmol CO_2_m^-2^s^-1^±0.08 vs wet 11.5±0.80), higher chlorophyll absorbance (dry 36.2 SPAD±1.4 vs wet 29.8±3.2), greater stomatal conductance (dry 0.174 mol H_2_Om^-2^s^-1^±0.003 vs wet 0.08 ±0.005), higher WUE (dry 7.29 µmol CO_2_/H_2_Om^-2^s^-1^±0.8 vs wet 1.40±0.2), greater nitrogen concentration (dry 1.80±0.20% vs wet 1.39±0.18%) and thicker leaves (dry 0.19 mm± 0.03 vs wet 0.17±0.04; all p<0.05; Figures 3a, S3). Further, the dry ecotype compared to the wet ecotype had significantly lower transpiration rates (dry 2.12 µmol H_2_Om^-2^s^-1^±0.2 vs wet 4.46±0.02), bolted earlier by 23 days (dry JD=219.9±3.6 vs wet 243.1±5.0) and flowered earlier by 16 days (dry JD=246.0±4.0 vs wet 262.3±4.2; Figures 3b, S3). This combination of traits indicate adaptation to dry conditions. Similar to the driest site, the wet ecotype compared to the dry ecotype was taller by ∼20 cm (wet 71.1 cm±16.9 vs dry 54.9±3.7) and had greater diameter by ∼40 cm (wet 91.3 cm±18.8 vs dry 50.3±4.0; all p<0.05; Figures 3c, S3).

##### Wet Site

Finally, at the wet site, the dry ecotype compared to the wet ecotype again demonstrated significantly greater photosynthetic rates (dry 15.0 µmol CO_2_m^-2^s^-1^±0.47 vs wet 12.3±0.29), higher transpiration rates (dry 3.78 µmol H_2_Om^-2^s^-1^±0.28 vs wet 2.36±0.25), and had thicker leaves (0.19 mm±0.02 vs 0.16±0.01; all p<0.05; Figures 3a, S3). Furthermore, the dry ecotype flowered earlier (dry JD = 219.0±3.5 vs wet 237.4±2.0) than the wet ecotype at the wet site (Figure 3b). The wet ecotype, at its homesite, compared to the dry ecotype was significantly taller by ∼100 cm (wet 160.2 cm±9.2 vs dry 59.9±15.9), had greater leaf width (wet 1.1 cm±0.3 vs dry 0.6±0.04), larger diameter (wet 107.1 cm±7.3 vs dry 47.3±6.2), greater total biomass by 200 g (wet 231.0 g±52.1 vs dry 27.6±4.9) and greater seed biomass (wet 9.7 g±3.2 vs dry 0.22±0.16; all p<0.05; Figures 3c, S3). This combination of traits indicate adaptation to wet conditions, where light and nutrients are limiting.

##### Rainout

Rainout treatments revealed ecotype-specific response to drought, with the wet ecotype generally exhibiting the strongest negative response. At the dry site, under rainouts compared to ambient, the wet ecotype had a significant reduction in photosynthetic rate (7.1 μmol CO_2_m^-2^s^-1^±0.9 lower), height (15.6 cm±1.3 lower), total biomass (18.2 g±2.4 lower), and diameter (12.4 cm±3.1 lower; Figures 3a-b, S3). In contrast, the dry ecotype showed significantly higher WUE (1.2 μmol CO_2_/H_2_Om^-2^s^-1^±0.9 increase) under rainouts relative to ambient and significantly lower transpiration rates (Figure S3), consistent with drought tolerance. Rainouts delayed flowering of all ecotypes by 5-15 days at the dry site (Figure 3c).

Finally, at the wet site, rainouts delayed flowering of all ecotypes by 6-17 days (Figure 3c). The wet ecotype demonstrated significantly reduced photosynthetic rates (4.2 μmol CO_2_±0.4 decline), WUE (2.3 μmol CO_2_/μmH_2_O±0.09 decline), height (35.3 cm±0.9 decline), seed biomass (4.8 g±0.8 decline), and stomatal conductance (0.25 mol H_2_O±0.07 decline; Figures 3, S3; all p<0.05) under rainouts compared to ambient. In contrast, under rainouts the dry ecotype showed significantly higher chlorophyll absorbance (5.2 SPAD±0.4 increase), photosynthetic rates (2.7 μmol CO_2_±0.7 increase), stomatal conductance (0.11 mol H_2_O±0.4 increase), WUE (1.9 μmol H_2_O±0.2 increase), height (29 cm±1.4 increase), leaf width (0.19 cm±0.05 increase), and nitrogen concentration (0.46%±0.11 increase; Figures 3, S3; all p<0.05).

#### 3.1.3. Principal Components Analysis (PCA) of Functional Traits

Multivariate trait associations revealed clear ecotype differentiation and environmental clustering across sites and rainout treatment. In nearly all PCAs analyzed, axis 1 explained over 50% of the total variance, with axis 2 accounting for 15–20%. Consequently, our interpretation focuses on axis 1. In the PCA by ecotype, axis 1 was significantly correlated with mean annual precipitation across all three ecotypes (p<0.05; Figure S4), highlighting a rainfall-trait relationship consistent with local adaptation.

##### By Ecotype

Trait coordination differed among ecotypes: for the dry ecotype (Figure 4a), axis 1 was strongly associated with WUE and leaf thickness—traits aligned with a drought tolerance strategy. In contrast, for the wet ecotype (Figure 4b), most traits corresponded to a growth/ competition strategy such as height and total biomass.

**Figure 4.**
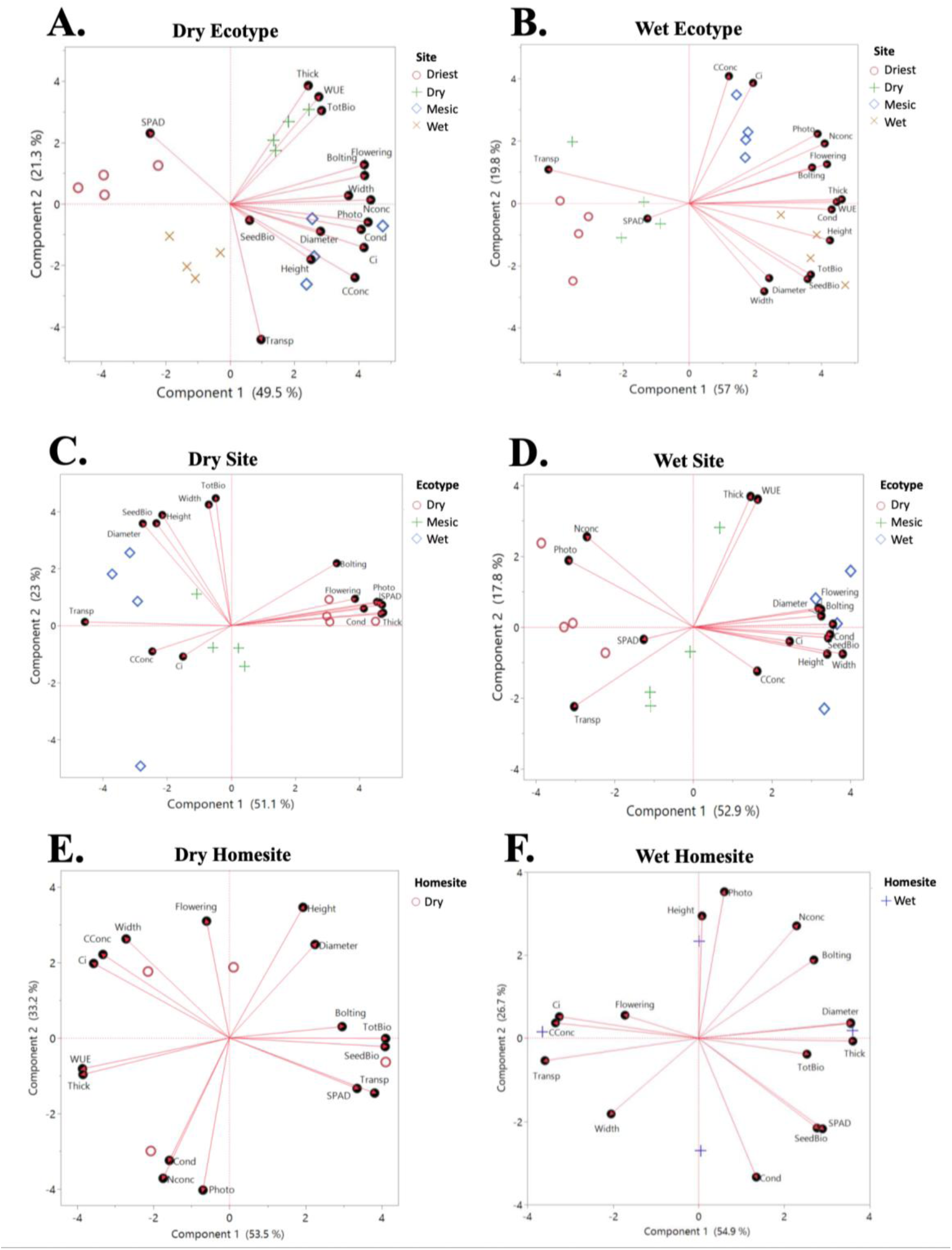
A. Principal component analysis (PCA) biplot of trait data by ecotype. A. dry ecotype and B. wet ecotype (mesic in the supplement) where colors and shapes indicate site (driest, dry, mesic, and wet). A PCA biplot of trait by site: C. dry site and D wet site in peak season (mesic and driest site in the supplement) where colors and shapes indicate ecotype (dry, mesic, or wet). A PCA biplot of trait by homesite. E. dry homesite and F. wet homesite in peak season where colors and shapes indicate ecotype (dry or wet). Lines indicate plant traits, and points are the means.

##### By Site

The PCA analyzed by site demonstrated trait coordination among sites along axis 1. At the dry (Figure 4c) and driest sites (Figure S5), increased WUE, greater leaf thickness, and increased chlorophyll absorbance loaded strongly on axis 1 and were associated with the dry ecotype. At the wet site (Figure 4d), nearly all morphological traits were associated with the wet ecotype. These patterns indicate that site-level environmental conditions shape trait integration and ecotype performance.

##### Homesite

The PCA by homesite also revealed strong evidence for local adaptation through trait coordination. The dry homesite (Figure 4e) showed positive loading of total biomass, seed biomass, transpiration rate, SPAD, and bolting with axis 1, while WUE and leaf thickness were negatively associated. At the wet homesite (Figure 4f), canopy diameter, leaf thickness, and total biomass were positively associated with axis 1, whereas internal CO₂ concentration, carbon concentration, transpiration rate, and flowering were negatively associated. These patterns of trait coordination at homesites support local adaptation, with each ecotype exhibiting trait combinations optimized for performance at their homesite.

##### Rainouts

Comparing PCA under rainouts, we identified specific traits linked to each ecotype. For example, under rainouts, leaf thickness, WUE, bolting, flowering, photosynthetic rate, and nitrogen concentration were linked to the dry ecotype (Figure S6a). However, in ambient conditions, canopy diameter, height, total biomass, carbon concentration, and leaf width were associated with the wet ecotype (Figure S6b).

### 3.2. Objective 2: Gene Expression Analyses

#### Gene Expression Characteristics

Here we focus on a subset of plants for differential gene expression analysis: the dry and wet ecotypes, driest, dry, and wet sites, and treatment (ambient and rainout). In the differential gene expression analysis, we identified 77,471 genes among these samples with each sample with a FastQC score of 97.5% or higher. Of the identified genes, 66,037 genes had annotations. A list of candidate genes identified by ecotype, site, and rainout treatment is presented in Table S5.

#### Ecotype

We identified 11,278 differentially expressed genes (DEGs) with a main effect of ecotype (Figure 5a, Table S3). In the dry ecotype, GO terms were significantly enriched for root epidermal cell, root differentiation, chlorophyll metabolism, and regulation of stomatal closure (Figure S7a; Table S4). In the wet ecotype, GO terms were significantly enriched in genes related to gibberellin response, trichome differentiation, and auxin-activated pathway (Figure S7a; Table S4). Taken together, this indicates a strategy of water conservation and belowground investment for the dry ecotype and growth promotion and aboveground development for the wet ecotype.

**Figure 5.**
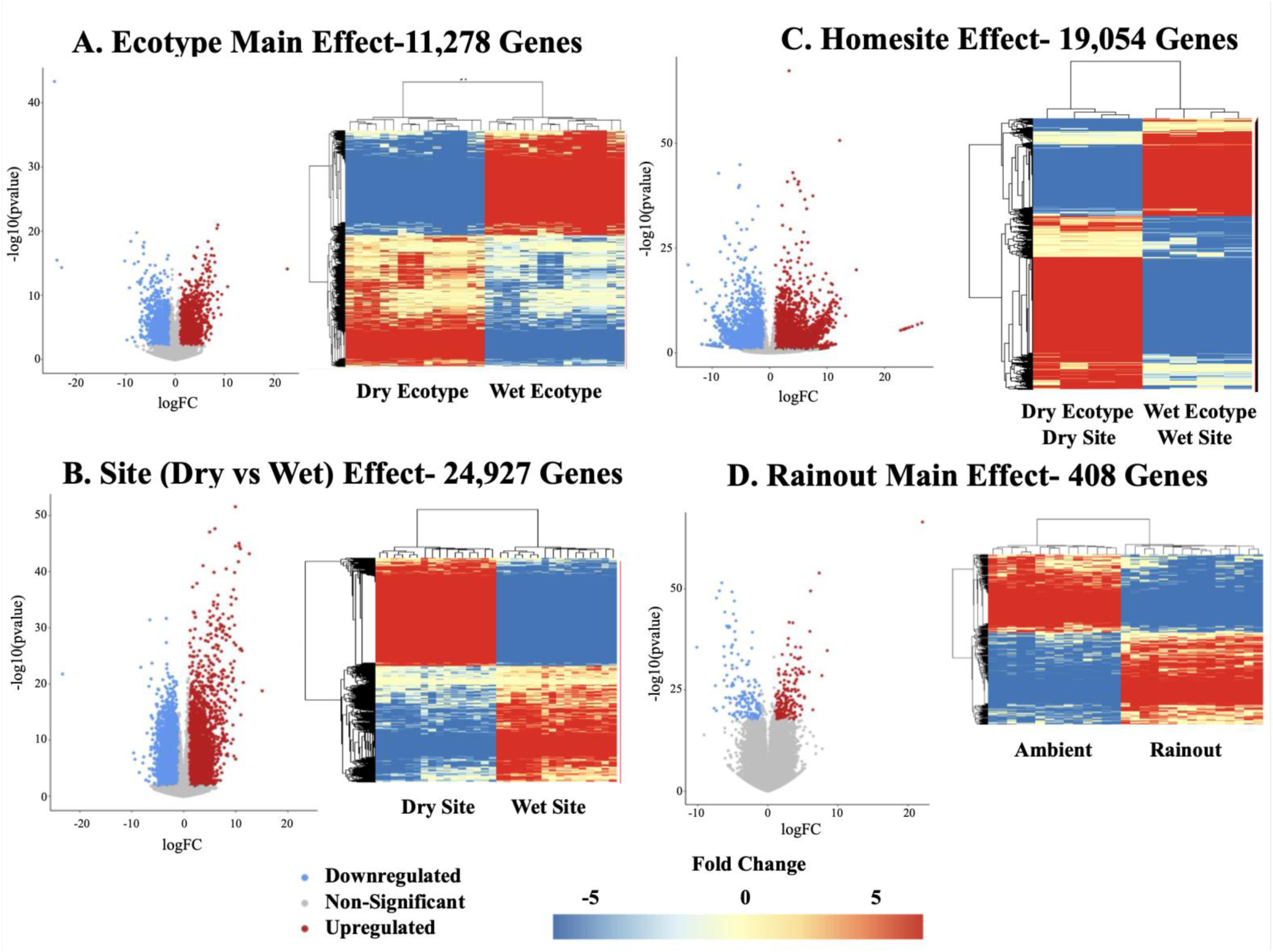
Heatmaps and volcano plots depicting the following: A. 11,278 differentially expressed genes between the wet and dry ecotypes, sites combined (ecotype main effect). B. 24,927 differentially expressed genes between the wet and dry sites, ecotypes combined (site main effect) C. 19,054 differentially expressed genes between the wet and dry ecotypes in their respective home sites (homesite effect). D. 408 differentially expressed genes between ambient and rainout conditions, ecotypes and sites combined (rainout main effect). Red points indicate upregulated genes; blue points indicate downregulated genes in volcano plots with unsignificant genes (p adj > 0.05) are indicated by gray points. The fold change in heatmaps indicated by blue (negative expression) or red (positive expression).

#### Site

Site comparisons identified 24,927 DEGs between dry and wet sites (Figure 5b, Table S3). At the dry site, GO terms were significantly enriched for root cap of primary root, response to stress, defense response, and lateral root tip (Table S4; Figure S7b). In contrast, at the wet site, GO terms were significantly enriched in growth-related processes such as biosynthetic and metabolic processes and auxin response shoot system meristem, response to auxin, chloroplast organization, and reproductive bud (Table S4; Figure S7b). Comparing the wet and the driest site, we found 10,015 DEGs (Table S4), showing enrichment for GO terms related to defense response and abscisic acid transport at the driest site (Table S4). Taken together, this indicates a strategy of stress tolerance and root-driven resource acquisition for the dry site and growth and reproductive development for the wet site.

#### Homesite

Most informative are comparisons of gene expression between ecotypes at their homesites revealing genetic signatures of local adaptation. Specifically, comparing the dry ecotype at the dry site and wet ecotype at the wet site, we identified 19,054 DEGs (Figure 5c). In the wet homesite, 8,713 genes were significantly enriched for GO terms related to seed maturation, gibberellin mediated signaling pathways, shoot system morphogenesis, and apical meristem (Figure S7c; Table S4). In the dry homesite, 10,341 genes were significantly enriched for ABA signaling, photosynthesis, root epidermis, and pollen development (Figure S7c; Table S7). In summary, this indicates a strategy of stress-adaptation at the dry homesite and growth-oriented development at the wet homesite, consistent with local adaptation to contrasting environments.

#### Rainout

Gene expression patterns reflected distinct responses to rainout and ambient treatment (across sites and ecotypes) with 407 DEGs showing a treatment effect (Figure 5d, Table S3). Under rainouts, GO terms were significantly enriched for root apical meristem, cellular water homeostasis, heat shock protein signaling, and chloroplast modification (Figure S7d; Table S4). In ambient conditions, GO terms were significantly enriched for vascular growth, stem epidermis, first order inflorescence axis, and regulation of growth (Figure S7; Table S4). This indicates a strategy of stress mitigation under experimental drought, and developmental progression and structural investment under ambient conditions.

### 3.3. Objective 3: Linking Gene Expression to Phenotype with Co-expression Networks Ecotype

To uncover the molecular basis of adaptive divergence between ecotypes, we analyzed gene co-expression networks linked to phenotypic traits. These networks reveal a striking contrast in adaptive strategies between wet and dry ecotypes that may drive local adaptation. First, in the dry ecotype (Figures 6a, S8a), we identified upregulated genes involved in stress responses, photosynthesis, and gene regulation (notably, *chlorophyll A-B binding, drought-induced-19, chloroplast drought-induced protein,* Cluster 26; *DREB2A,* Cluster 13) which were co-expressed with transpiration rate, internal carbon dioxide concentration, and height (red stars, Figure 6a). Upregulated genes related to growth (e.g., *ARF-16,* Cluster 33) were co-expressed with growth rate and transpiration rate (Figure 6a; Table S6). These patterns reveal that the dry ecotype coordinates gene expression around stress-response pathways, reflecting a strategy prioritizing resource conservation in challenging environments.

**Figure 6.**
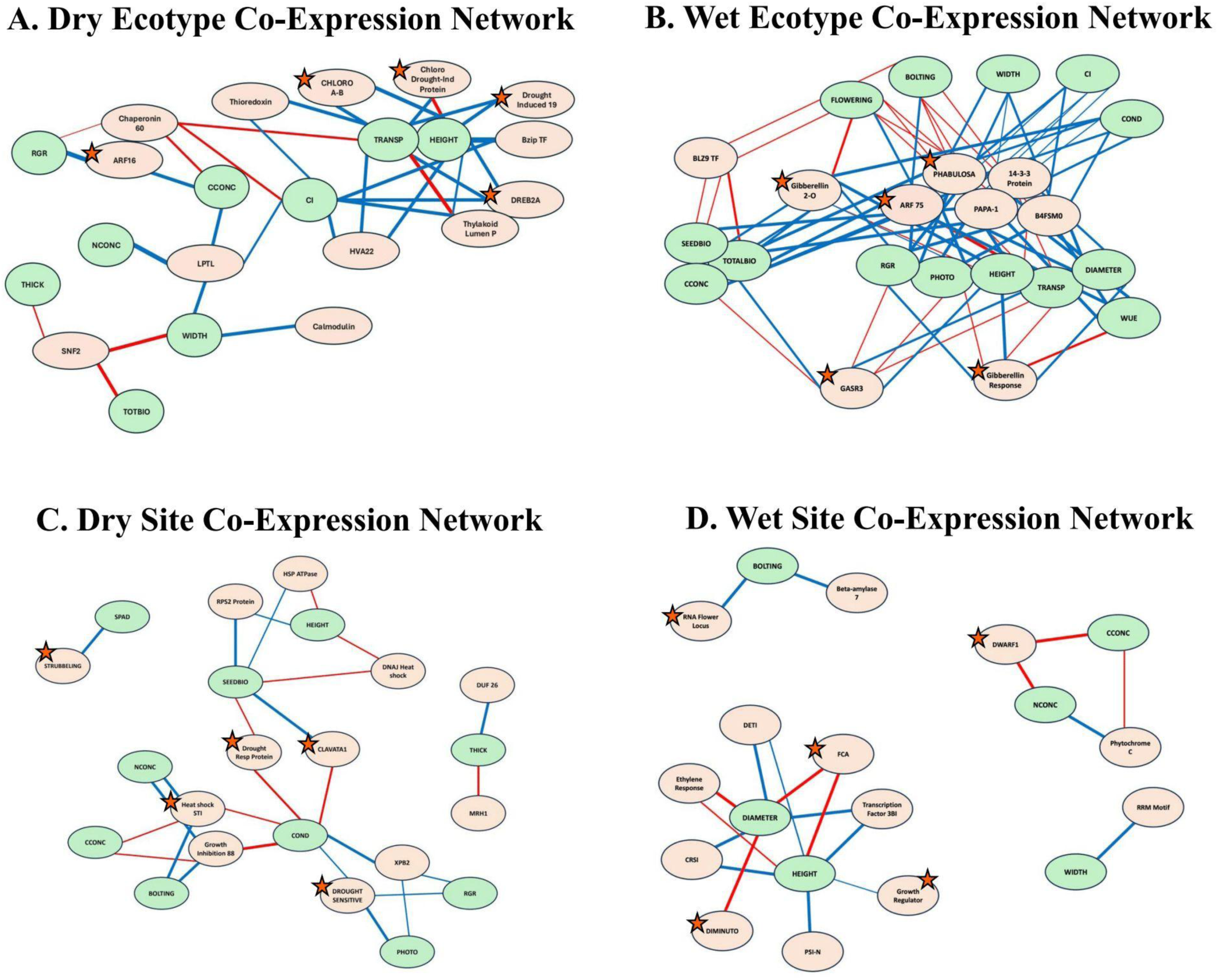
Co-expression network analysis of gene-trait associations. A. dry ecotype, B. wet ecotype, C. dry site, and D. wet site. Candidate genes were selected based on significant gene clusters from WGCNA with more than 3-fold change in expression and a known annotation. Green circles represent measured traits, while orange circles denote genes. Stars indicate genes with a well-documented function and provide insight into adaptive mechanisms. Line thickness corresponds to the strength of the association, with red lines indicating positive correlations and blue lines representing negative correlations. This network visualization highlights key relationships between gene expression and phenotypic traits, providing insights into potential functional connections underlying trait variation.

In contrast, in the wet ecotype, we identified genes linked to ten traits (Figure S8b; Table S6) such as flowering, stomatal conductance, height, and total biomass, co-expressed with upregulated genes related to auxin and growth (e.g., *ARF-75, PHABULOSA*, Cluster 11; Figure 6b; Table S6). Furthermore, upregulated genes involved in response to the growth hormone gibberellin (*gibberellin 2-O*, Cluster 30; *gibberellin response*; *GASR3* Cluster 29; Figure 6b; Table S6) that were co-expressed with nine traits including height and growth rate. This network suggests that the wet ecotype’s adaptive strategy emphasizes growth and biomass accumulation in favorable conditions.

#### Site

Site-specific gene–trait associations underscore how local environment shape both traits and gene expression. At the dry site, genes involved in osmotic regulation (e.g., *DROUGHT SENSITIVE 1,* Cluster 5; Figures 6c, S8c; Table S6) were co-expressed with stomatal conductance, growth rate, and photosynthetic rate. Genes involved in floral meristem regulation (e.g., *CLAVATA1,* Cluster 6) and stress response (e.g., *drought-responsive protein*, Cluster 6; Figure 6c) were co-expressed with seed biomass and stomatal conductance. Heat-shock-related genes (e.g., *Heat-Shock STI,* Cluster 4) were positively linked to nitrogen concentration, carbon concentration, and bolting. Finally, genes involved in root growth (e.g., *STRUBBELING-RECEPTOR FAMILY 7*, Cluster 18), was co-expressed with chlorophyll absorbance (Figures 6c, S8c; Table S6). These networks highlight how stressors at the dry site selectively upregulate genes related to stress tolerance.

In contrast, at the wet site, genes involved in regulating flowering (e.g., *FCA*, Cluster 8) and those involved in growth regulation (e.g., *Growth Regulator*, *DIMINUTO,* Cluster 8) were co-expressed with height and diameter (Figures 6d, S8d; Table S6). Genes involved with flowering (e.g., *RNA flowering locus*, Cluster 10) were co-expressed with bolting. Finally, genes related to growth (e.g., *DWARF1:* Cluster 2) were co-expressed with nitrogen and carbon concentration (Table S6; Figure 6d). Co-expression networks at the wet site suggest that favorable conditions promote a strategy geared toward maximizing growth and development.

#### Homesite

Gene–trait associations of the ecotypes reflected local environmental conditions, supporting local adaptation. At the dry homesite, several genes involved in regulation of flowering (e.g., *FRIGIDA*, Cluster 16; Figures 7a; S8e) were co-expressed with six traits including total biomass and leaf width. Further, we identified downregulated genes involved in root growth (e.g., *root hair defective-5,* Cluster 10) were co-expressed with nine traits including transpiration rate and WUE (Figure 7a; Table S6). Other upregulated genes were involved in auxin response and photosynthesis (notably *auxin response-8, photosystem reaction center*, Cluster 9) were each linked to multiple traits including WUE, photosynthetic rate, and growth rate (Figure 7a; Table S6). Several genes involved in root formation (e.g., *aberrant root formation-P4*, Cluster 17) were linked to seed biomass, height, and stomatal conductance at the dry homesite (Figure 7a; Table S6). Our network reflects local adaptation of the dry ecotype to water-limited conditions, with coordinated regulation of genes that optimize water-use efficiency under low moisture.

**Figure 7.**
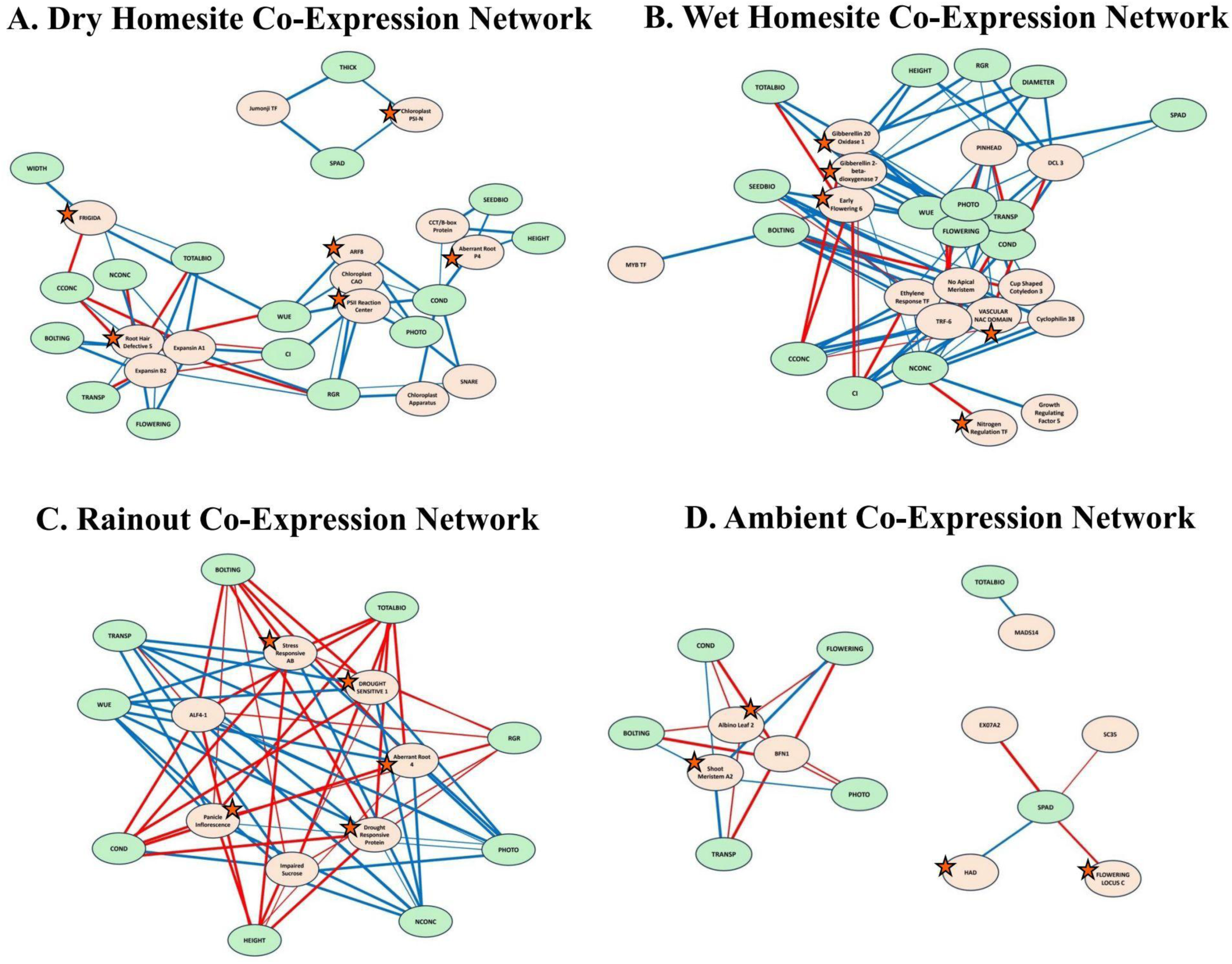
Co-expression network analysis of gene-trait associations. A. dry homesite and B. wet homesite, C. rainout, D. ambient treatments. Candidate genes were selected based on significant gene clusters from WGCNA with more than 3-fold change in expression and a known annotation. Green circles represent measured traits, while orange circles denote genes. Stars indicate genes with a well-documented function and provide insight into adaptive mechanisms. Line thickness corresponds to the strength of the association, with red lines indicating positive correlations and blue lines representing negative correlations. This network visualization highlights key relationships between gene expression and phenotypic traits, providing insights into potential functional connections underlying trait variation.

In comparison, at the wet homesite, many gene clusters were strongly associated with multiple traits (Figure S8f). For example, downregulated genes involved in nitrogen regulation (e.g., *Nitrogen Regulation Transcription Factor*, Cluster 26) were co-expressed with nitrogen concentration. Genes involved in vascular tissue generation (e.g., *VASCULAR-RELATED DOMAIN-6*, Cluster 23) were co-expressed with several traits including flowering, seed biomass, and photosynthetic rate (Figure 7b; Table S6). Additionally, gibberellin-related genes (e.g., *Gibberellin 20-Oxidase 1,* Cluster 5*, Gibberellin 2-beta-dioxygenase-7*, Cluster 6) and genes involved in flowering (e.g., *Early Flowering 6*, Cluster 23) were broadly co-expressed with most traits, including height, photosynthetic rate, and seed biomass (Figure 7b; Table S6). This network highlights local adaptation of the wet ecotype, with genes promoting growth and reproduction, reflecting optimization of development under abundant soil moisture.

#### Rainout Treatment

Of the 407 DEGs expressed under rainouts across sites and ecotypes, 146 were significantly correlated with a measured phenotype (Figure S8g-h). For example, under rainouts we identified several upregulated genes that are involved in root development and stress response (notably *Aberrant Root Formation-4, stress-responsive-AB, DROUGHT SENSITIVE-1, drought responsive protein,* and *Panicle Inflorescence*, Cluster 3). These genes were co-expressed with nine traits including WUE and bolting (Figure 7c; Table S6). In contrast, in ambient conditions (Figure 7d), genes involved in flowering (e.g., *FLOWERING LOCUS C,* Cluster 1) or nutrient uptake (e.g., *Haloacid dehalogenase* (*HAD*), Cluster 1) were co-expressed with chlorophyll absorbance (Figure 7d). Also in ambient conditions, genes involved in vascular tissue growth (e.g., *shoot apical meristem A2, Albino Leaf* 4, Cluster 3) were co-expressed with five traits including flowering, photosynthetic rate, and stomatal conductance. Taken together, our networks demonstrate gene–trait linkages associated with stress tolerance, highlighting adaptive responses to both natural rainfall gradients and experimental drought conditions.

## 4. Discussion

This study sheds light on how form, function and growth of plant ecotypes are influenced by rainfall gradients and experimentally reduced rainfall, by linking gene expression and plant functional traits to characterize local adaptation (summarized in Figure 8). Here, we explore how local adaptation is indicated by plant traits indicating coordination of trait strategies (Grime, 1988): stress tolerant traits in the dry ecotype in response to water limitation and competitive growth traits (Jardine et al., 2021) in the wet ecotype due to light and nutrient limitation. We then discuss the gene expression differences between ecotypes, which at the genetic level, support the divergent adaptive phenotypes. Further, using the experimental rainfall manipulation, we show that rainfall is the driving selective force leading to divergence of these ecotypes. Finally, by connecting gene expression to traits like photosynthetic rate, biomass, and height, we reveal the genetic basis of adaptive phenotypes as related to rainfall. This integrative approach offers a unique perspective on the genetic basis of ecological performance in natural plant communities (Johnson et al., 2022) under changing climatic conditions (Benning et al., 2023).

**Figure 8.**
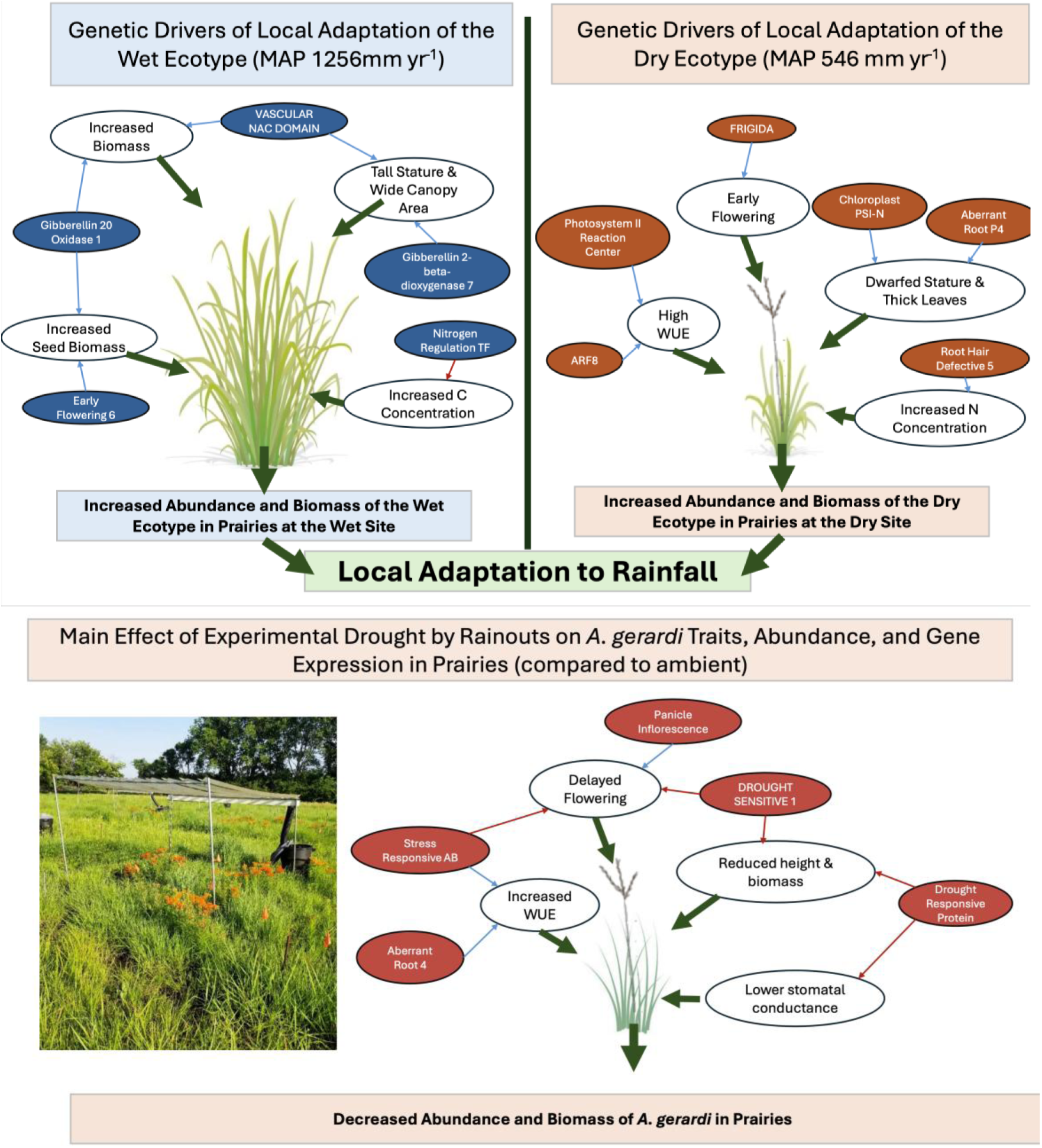
This model illustrates the adaptive strategies of wet and dry ecotypes in response to their respective home sites. In the wet ecotype, upregulated genes related to growth regulation, gibberellin signaling, and biomass accumulation are associated with increased resource capture and competitive success under favorable moisture conditions. In contrast, the dry ecotype shows enhanced expression of drought-tolerant genes involved in water retention, photosynthetic efficiency, and stress response mechanisms, enabling survival in arid environments. The lower panel illustrates the influence of experimental drought by rainouts on plant performance, trait variation, and gene expression. Our conceptual model highlights the role of gene-environment interactions in local adaptation and the importance of assessing both trait and transcriptomic data to understand the molecular basis of ecological success in plant communities.

### 4.1. Evidence of plot-level local adaptation to rainfall linked to functional traits

We found the highest cover and biomass of the wet and dry ecotypes at the plot-level in their home site and lower at foreign sites, supporting local adaptation of *A. gerardi* (Galliart et al., 2019; 2020). It is well-established that climate drives intraspecific variation and local adaptation (Turesson, 1922; McMillan, 1965; Savolainen et al., 2013) where numerous studies have demonstrated that populations often exhibit higher fitness in their home environments, reflecting patterns of local adaptation (Joshi et al., 2001; Leimu & Fischer, 2008; Lortie & Hierro, 2022). However, few studies integrate local adaptation with transcriptomics and traits across environmental gradients, especially in native, non-model species in ecological communities (Johnson et al., 2022; Sytsma et al., accepted). We address this gap by integrating ecological and genomic approaches to reveal mechanisms of local adaptation in *A. gerardi* across a natural rainfall gradient and in response to experimental drought (Figure 8) in grassland communities. Since rainfall putatively drives ecotypic differentiation (Gray et al., 2014), our use of cross-transplantation in community settings enabled realistic detection of local adaptation.

Our findings reveal clear physiological and morphological divergence and coordinated trait strategies among ecotypes that underpin local adaptation to the rainfall gradient. The dry ecotype exhibited classic drought-adaptive traits, including higher photosynthetic rates, improved water-use efficiency, thicker leaves, and elevated leaf nitrogen despite smaller stature (Grime, 1988; Jardine et al., 2021; Ochola et al., 2024). Its earlier flowering suggests a drought escape strategy at the end of the growing season or adaptation to shorter growing seasons (Shavrukov et al., 2017), consistent with patterns seen in arid-adapted species showing early phenology, reduced size, and efficient water use (Castillioni et al., 2022). In contrast, the wet ecotype prioritized growth, producing wider leaves and more biomass, and growing taller particularly at the wet site. This reflects a broader trend in which ecotypes from wetter regions favor rapid growth and resource acquisition (Jardine et al., 2021). These differences illustrate trade-offs between growth and stress tolerance (Grime, 1988), providing mechanistic evidence of local adaptation through trait divergence (Figure 8).

Rainout manipulations demonstrated that rainfall is the selective pressure driving divergent ecotypes, further corroborating support for regional climate ecotypes and local adaptation. Rainfall imposes strong selective pressure in many organisms, often driving population differentiation in traits related to growth, reproduction, and survival (Siepielski et al., 2017). Consistent with this, rainouts delayed flowering in all ecotypes, especially at the dry site, linking water availability to reproductive timing in *A. gerardi* (Dietrich & Smith, 2016). In general, rainouts reduced the performance of all ecotypes but disproportionately impacted the wet ecotype, underscoring its adaptation to high-moisture environments and reduced drought tolerance.

At the wet site, we found additional strong evidence of rainfall as a selective pressure. Interestingly, under rainouts, the dry ecotype’s biomass, cover, photosynthetic rate, and height *increased* compared to ambient, suggesting that reduced rainfall in wetter environments may favor drought-adapted ecotypes (Metz & Tielborger, 2023). Additionally, under rainouts at the dry site, the wet ecotype showed reduced WUE compared to ambient, while the dry ecotype showed increased WUE, indicating greater drought tolerance (Tardieu, 2022). These results demonstrate that rainfall not only drives local adaptation but may also shapes functional trait divergence in ecological communities.

### 4.2. Gene expression profiles reflect divergent adaptive phenotypes

Distinct patterns of gene expression were correlated between ecotypes, sites, and rainfall treatments, suggesting local genetic adaptation (Savolainen et al., 2013; Hoffman et al., 2018; Figure 8). These findings suggest that key physiological pathways were selected in each ecotype to optimize their physiological responses to local environmental conditions.

First considering ecotype, the wet ecotype upregulated genes involved in carbon gain and growth, reflecting an adaptive strategy favoring resource acquisition and growth investment in high-moisture environments. Building on findings by Galliart et al. (2020), our data also show further upregulation of gibberellic acid pathways (*gibberellin 2-O*, *GASR3*)—key to germination, shoot elongation, and reproductive transitions (Shah et al., 2023)—likely contributing to the wet ecotype’s greater height and biomass. In contrast, the dry ecotype upregulated genes related to root development (*Chaperonin 60*), dehydration response (*DREB2A)*and stomatal complex formation (*chloroplast drought-induced protein*), aligning with drought-adaptive strategies aimed at enhancing water uptake and regulating gas exchange (Pardo et al., 2020; Gupta et al., 2024; Zhang et al., 2025b). These contrasting gene expression profiles reflect distinct physiological strategies and reinforce the role of local adaptation at the genetic level (Figure 8).

The impact of precipitation on gene expression is further evident when comparing sites, with ecotypes combined, with contrasting wet and dry rainfall patterns. First, at the wet site, genes involved in growth (*Growth Regulator*, *DIMINUTO*) and flowering (*FCA, RNA Flowering Locus*) were upregulated, reflecting an earlier onset of the growing season. In contrast, at the dry site, we found enrichment of genes involved in abscisic acid signaling (*ABI-3*), a key pathway in drought response (Aimar et al., 2014). These findings suggest that rainfall drives significant shifts in gene expression (Hamann et al., 2021), with the wet site promoting growth and reproductive pathways, while the dry site upregulates drought response mechanisms. Similarly, other studies (Lovell et al., 2016; Bhaskara et al., 2023) reported differential expression of drought-responsive genes across a precipitation gradient in *Panicum hallii*, supporting the idea that aridity selects for transcriptomic shifts associated with water conservation.

Considering gene expression of ecotypes in their homesite, our analysis revealed strong transcriptomic differentiation between wet and dry ecotypes where they have been strongly selected for by regional climate over thousands of years (Axelrod, 1985). At the wet homesite, enrichment of genes related to gibberellin signaling (*Gibberellin 20-Oxidase 1, Gibberellin 2-beta-dioxygenase 7*) suggests an environment-driven developmental shift favoring growth, resource acquisition, and competitive ability in resource-rich environments (Shah et al., 2023). Conversely, in the dry homesite, gene enrichment in processes related to early flowering (*FRIGIDA*), gas exchange (*photosystem reaction center),* and root growth (*root hair defective-5, aberrant root formation-P4)* indicates an adaptive focus on survival and resource conservation under water-limited conditions (Cardoso et al., 2015). Similar patterns have been observed in *Panicum* spp., where wet populations exhibit enrichment of genes involved in growth, while dry populations express genes associated with stress tolerance (Meyer et al., 2014; Lovell et al., 2018; Heckman et al., 2025). Together, homesite comparisons uncover expression patterns that show local adaptation is driven by transcriptome patterns leading to divergent physiological strategies (Figure 8).

Finally, under experimental drought, the enrichment of genes related to stress response (*stress-responsive-AB, DROUGHT SENSITIVE-1)* and regulation of flowering (*Panicle Inflorescence* ) suggests a coordinated developmental response to water limitation. These shifts in gene expression support a strategy of water balance, reduced growth, and reproduction under stress (Wu et al., 2022). Similar patterns in other species show enrichment in the same pathways during drought, highlighting convergent adaptive responses to water limitation (Lovell et al., 2016; Shrestha et al., 2023). Together, these results underscore how ecotypes cope with drought by modulating development and phenology to enhance drought resilience (Napier et al., 2023).

### 4.3. Co-expression networks link genes to traits and highlight the role of rainfall

Our co-expression networks connect gene expression to phenotypic traits, providing insight into the genetic architecture behind local adaptation (Figure 8). Specifically, co-expression networks connect transcriptomic profiles with traits and are instrumental in understanding plant biology and informing restoration efforts (Aoki et al., 2007). These findings underscore the importance of exploring how gene expression patterns influence adaptive traits, offering a deeper insight into the mechanisms that drive local adaptation (Savolainen et al., 2013; Kang et al., 2023).

Ecotype-specific patterns of gene expression and trait correlations reveal distinct adaptive strategies. In the dry ecotype, genes involved in photosynthesis and stress tolerance (e.g., *ARF-16, drought-induced-19, DREB2A;* Wu et al., 2022; Zhang et al., 2025b) were correlated with traits such as transpiration rate and height, pointing to regulatory networks that enhance water-use efficiency and growth in arid conditions. These findings are consistent with drought-adaptive strategies observed in *Brachypodium* sp. (Verelst et al., 2013; Wang et al., 2025). In contrast, the wet ecotype upregulated genes involved in gibberellin response and growth (*gibberellin response*, *GASR3, ARF-75;* Shah et al., 2023; Zheng et al., 2023) which were linked to competitive traits (Grime, 1988) including height and biomass. These ecotype-specific networks highlight distinct strategies—drought resilience in the dry ecotype versus growth in wet ecotype—offering insights into local adaptation and potential responses to changing environmental conditions.

Looking at the site differences, at the dry site, several genes were correlated with functional traits across ecotypes. Notably, genes known to be involved in drought tolerance (*drought-responsive protein, DROUGHT SENSITIVE 1,* and *Heat-Shock STI*; Soma et al., 2021; Ni et al., 2021) were linked to stomatal conductance and growth rate, suggesting that stress-response genes can influence growth under water stress. Similar patterns have been observed in *Arabidopsis*, where genes related to drought tolerance regulate growth and reproduction under water-limited conditions (Alam et al., 2022). At the wet site, genes associated with growth regulation and flowering control (*Growth Regulating Factor* and *FCA;* Wang et al., 2020), were co-expressed with traits linked with height and bolting. These site-specific gene-trait correlations highlight how environmental conditions shape ecotypic adaptations, supporting the genetic basis of local adaptation (Figure 8).

Considering gene expression differences at homesites (wet and dry), the clear differentiation observed between co-expression networks of the local ecotypes is particularly striking. At the wet homesite, genes related to growth (*VASCULAR-RELATED NAC DOMAIN-6*; Wang et al., 2022), were co-expressed with many adaptive traits including flowering and photosynthetic rate. Notably, genes involved in gibberellin response *(Gibberellin 20-Oxidase 1, Gibberellin 2-beta-dioxygenase 7)* were correlated with nearly all traits, emphasizing the importance of gibberellins in promoting growth in wet environments (Shah et al., 2023). In contrast, at the dry homesite, genes (*FRIGIDA*, *auxin response-8;* Hrmova & Mussain, 2021; Xu et al., 2022) were correlated with numerous traits including flowering, total biomass, and leaf width confirming results found in other species (Heckman et al., 2025). The contrasting co-expression networks provide strong evidence of how local conditions shape both gene expression and trait variation, supporting the role of local adaptation in plant survival.

Under rainouts, genes like *drought responsive protein* (Ying et al., 2023), *stress-responsive-AB* (Ji et al., 2016), and *DROUGHT SENSITIVE-1* (Abdel-Ghany et al., 2020) were correlated with traits including biomass and photosynthetic rate under experimental drought, demonstrating their involvement in stress response. Similar patterns have been observed in other studies (Hayford et al., 2022; Bhaskara et al., 2023), where drought-responsive genes were correlated with key traits such as biomass production and photosynthetic rate under drought. These co-expression patterns under rainouts highlight core stress-response genes as key to adaptation to rainfall (Figure 8), offering molecular targets for enhancing drought resilience in grasslands.

## 5. Conclusions

Our results demonstrate the value of linking gene expression to traits to uncover mechanisms of local adaptation under increasing drought. As a dominant foundation species, *Andropogon gerardi* shapes grassland community structure and function (Gibson, 2009; McCain et al., 2010; Ren et al., 2024), making its adaptation critical for predicting grassland responses to rainfall. By integrating trait and gene expression data, we identified genes associated with drought response and competitive ability—key fitness strategies—revealing molecular trade-offs between stress tolerance and growth. These insights provide targets for restoration, biofuels, and improving resilience in drought-prone systems (Zhang et al., 2015; Baer et al., 2019). One of the few ecological genomic studies conducted in long-term, realistic field settings (Johnson et al., 2022; Sytsma et al., accepted), our work highlights the need to understand genetic and phenotypic responses to rainfall to address challenges such as reduced productivity and forage in drought-prone ecosystems (Bao et al., 2019; Smith et al., 2020; Habte et al., 2022). Given that drought is a major driver of crop loss and reduced primary productivity (Dietz et al., 2021; Liu et al., 2023), our findings are timely and support more effective grassland restoration using climate-matched ecotypes.

## Author Contributions

**Jack Sytsma:** Writing- original draft, conceptualization, data curation, formal analysis, investigation, methodology, **Matthew Galliart:** Writing- review & editing, software, validation, **Kian Fogarty:** Writing- review & editing, data curation, investigation, **Kori Howe:** Writing- review & editing, investigation, **Bradley Olson:** Writing- review & editing, validation, software, **Sara Baer:** Writing- review & editing, resources, **David Gibson:** Writing- review & editing, resources, **Brian Maricle:** Writing- review & editing, resources, **Eli Hartung:** Writing- review & editing, investigation, **Loretta Johnson:** Writing- review & editing, conceptualization, funding acquisition, methodology, resources, project administration.

## Acknowledgements

Thanks to Z. Ren, D. Barfknecht, and N. Bishop for help in the field. Thanks to the National Center for Genome Resources and New Mexico State University INBRE for assisting in analysis and access to their server. Funding was provided by USDA Grant 2020-03665 and partially funded by the Kansas Academy of Science 2023 Student Research Grant, the 2023 Kansas Native Plant Society Bancroft Grant, and the U.S Department of Education Graduate Assistance in Areas of National Need Grant P200A240009 awarded to Sytsma.

## Conflicts of Interests

The authors declare that they have no competing interests.

## Data Accessibility

All data supporting the findings of this study will be available in a public repository upon publication. Raw sequencing reads, and processed genetic data will be deposited in the EMBL/GenBank repository, and trait measurements in a publicly accessible dryad repository Any additional data or materials can be made available upon reasonable request to the corresponding author.

## Benefit-Sharing Statement

This research was conducted within the United States on native plant species without involving genetic resources requiring prior informed consent under the Nagoya Protocol. Results will be shared through open-access publication, conference presentations, and outreach to land managers and restoration practitioners to support sustainable grassland management and conservation.

## Notes

### Competing Interest Statement

The authors have declared no competing interest.

