## Supplemental Data for "Taking genomics outdoors: linking local adaptation, trait variation, and gene expression in grass ecotypes across a rainfall gradient"

Supplemental Figure 1. Soil volumetric water content in each of the sites. A. driest, B. dry, C. mesic, and D. wet sites in 2022. Points indicate the mean of four blocks and error bars are the standard error. Ambient is indicated with blue and rainout treatment is indicated with red.

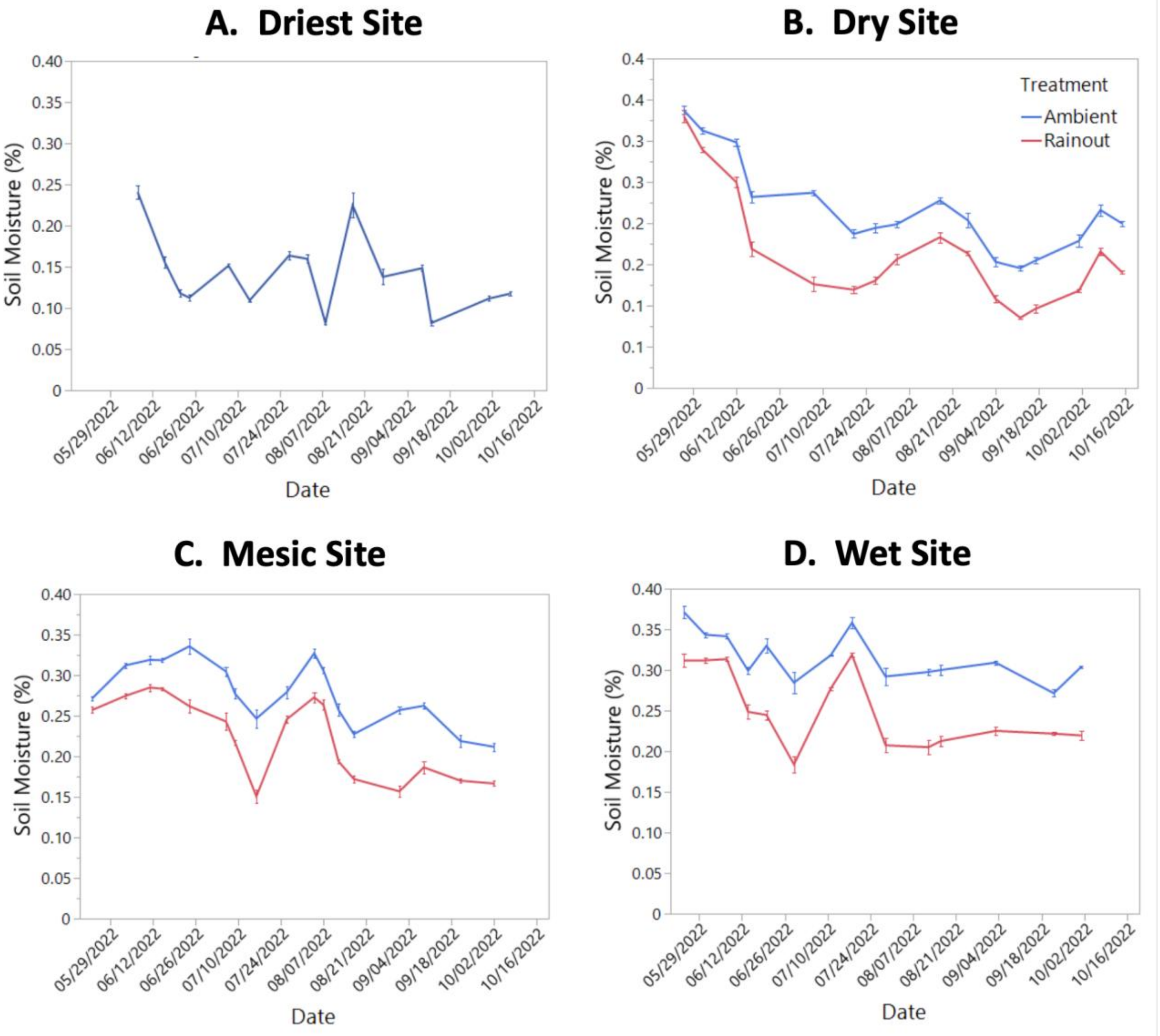

Supplemental Figure 2. Comparisons of community-level response variables. A. canopy cover and B. aboveground biomass in 2021. Ambient conditions are in blue, and rainout is in red. The points indicate the mean, error bars represent standard errors, and the different letters denote significantly differences between ecotypes and treatments within a site and year ($\alpha = 0.05$).

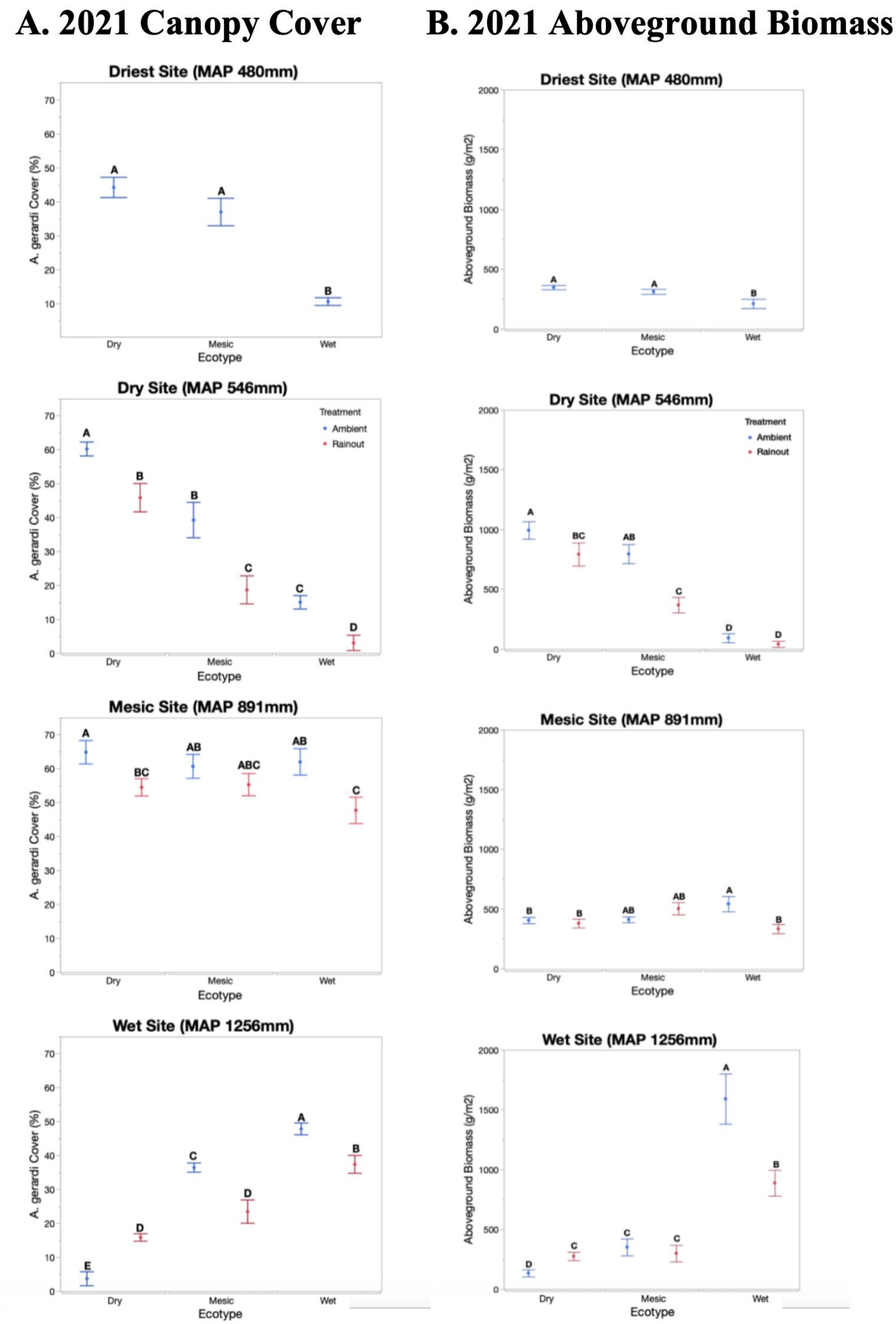

Supplemental Figure 3. Comparisons of traits. A. chlorophyll absorbance, B. stomatal conductance, C. transpiration rate, D. water use efficiency, E. blade width, F. seed biomass, G. total biomass, H. canopy diameter, I. leaf thickness, J. leaf nitrogen concentration, K. leaf carbon concentration, L. internal carbon dioxide concentration, and M. Julian date of bolting across sites (driest, dry, mesic, or wet) in peak season 2022. Ambient is blue, and rainout treatment is in red. The error bars represent standard errors, and the different letters denote significantly different groups among the comparisons within site ($\alpha = 0.05$).

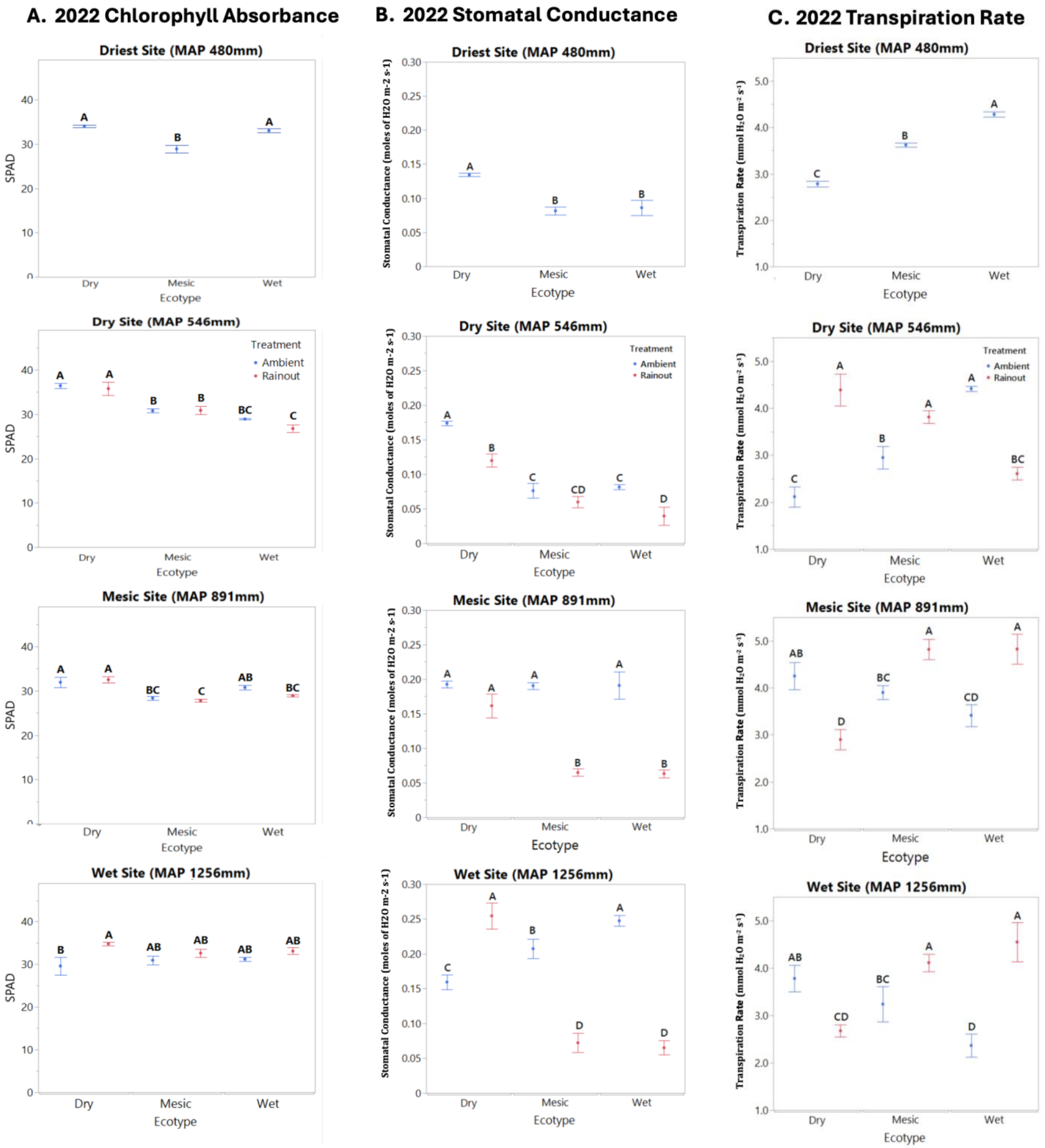

19

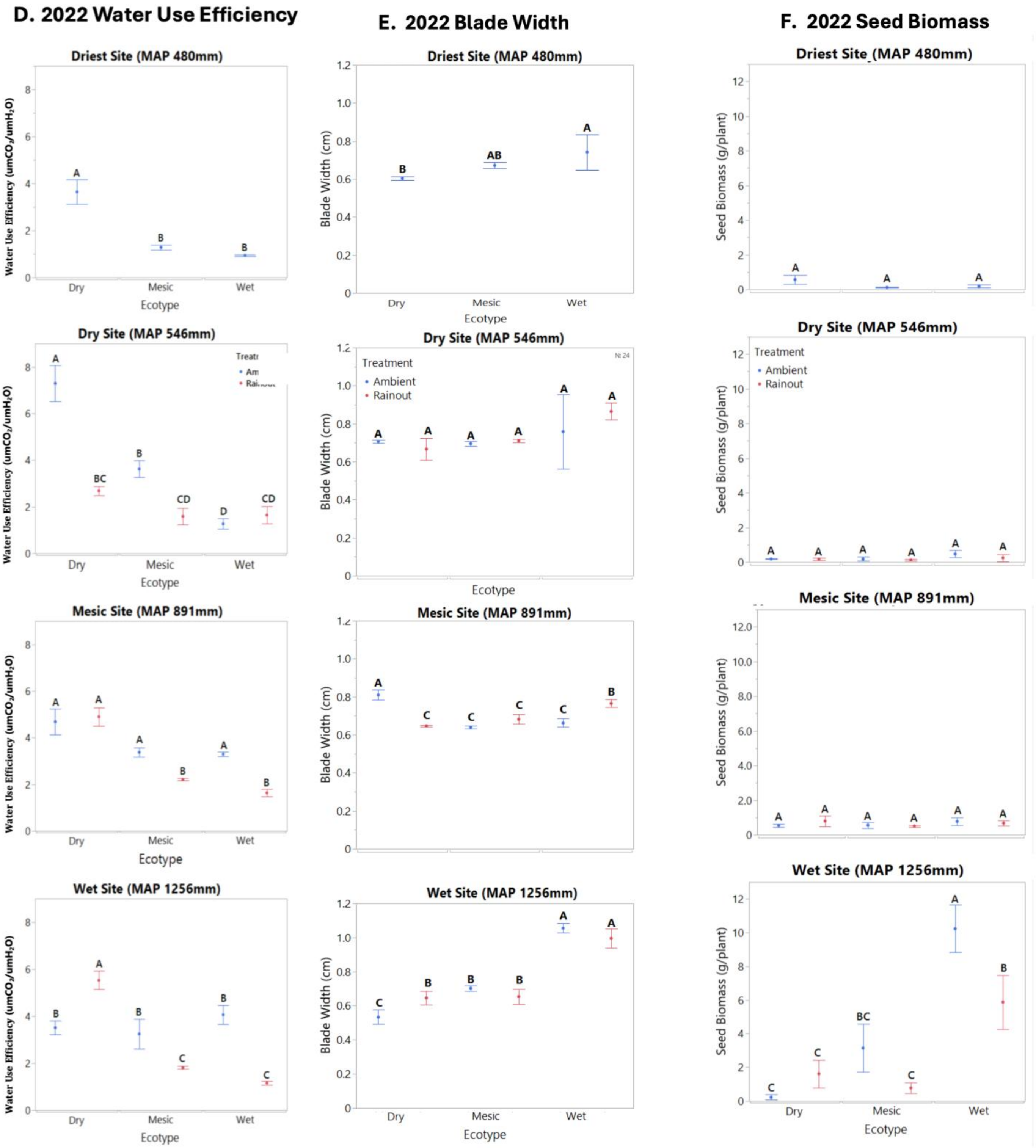
D. 2022 Water Use Efficiency
E. 2022 Blade Width
F. 2022 Seed Biomass
Driest Site (MAP 480mm)
Dry Site (MAP 546mm)
Mesic Site (MAP 891mm)
Wet Site (MAP 1256mm)
Water Use Efficiency (umCO2/umH2O)
Blade Width (cm)
Seed Biomass (g/plant)
Ecotype
Dry
Mesic
Wet
Treatment
Ambient
Rainout

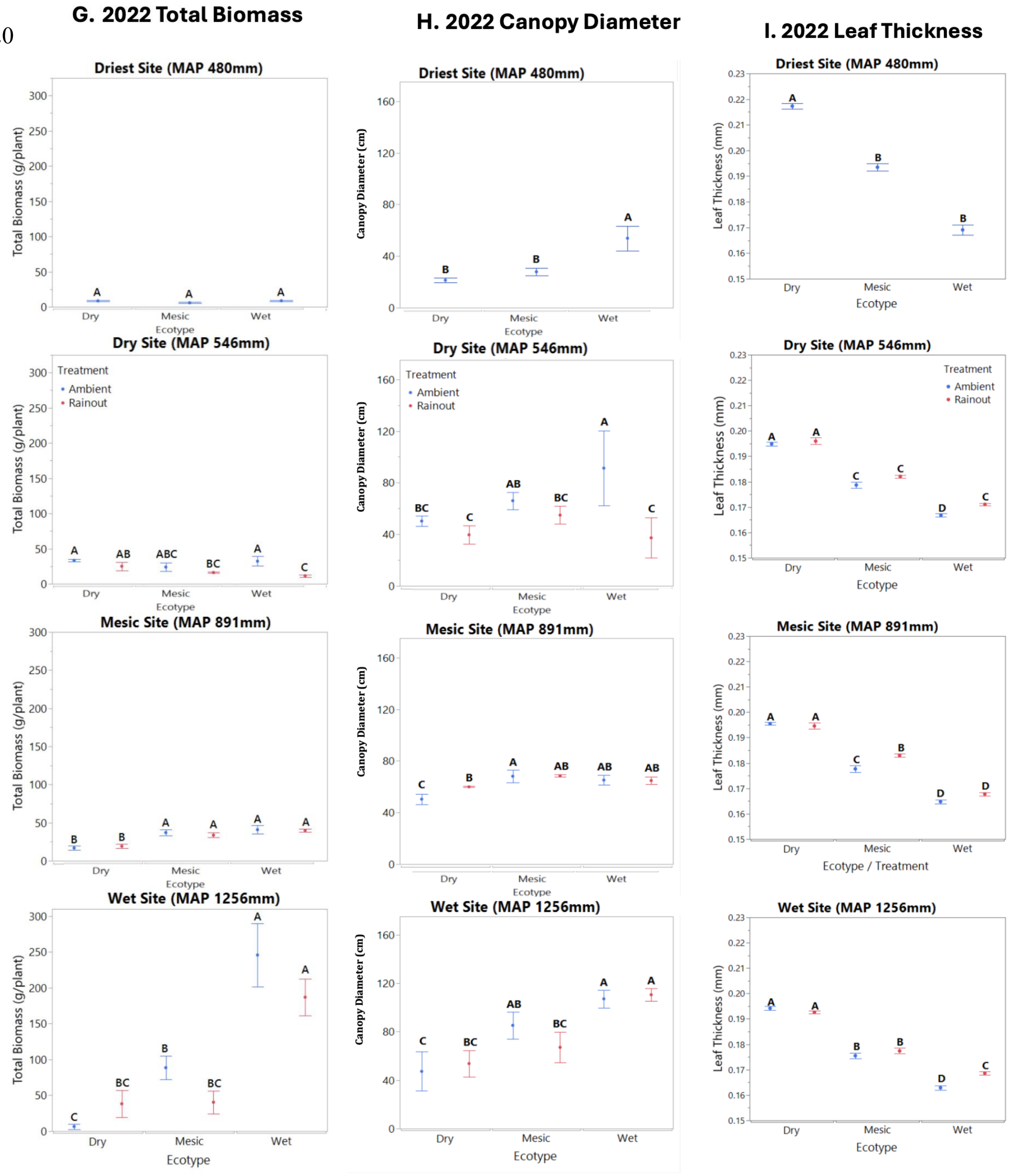
G. 2022 Total Biomass
H. 2022 Canopy Diameter
I. 2022 Leaf Thickness
Driest Site (MAP 480mm)
Dry Site (MAP 546mm)
Mesic Site (MAP 891mm)
Wet Site (MAP 1256mm)
Total Biomass (g/plant)
Canopy Diameter (cm)
Leaf Thickness (mm)
Ecotype
Ecotype / Treatment
Dry
Mesic
Wet
Treatment
Ambient
Rainout

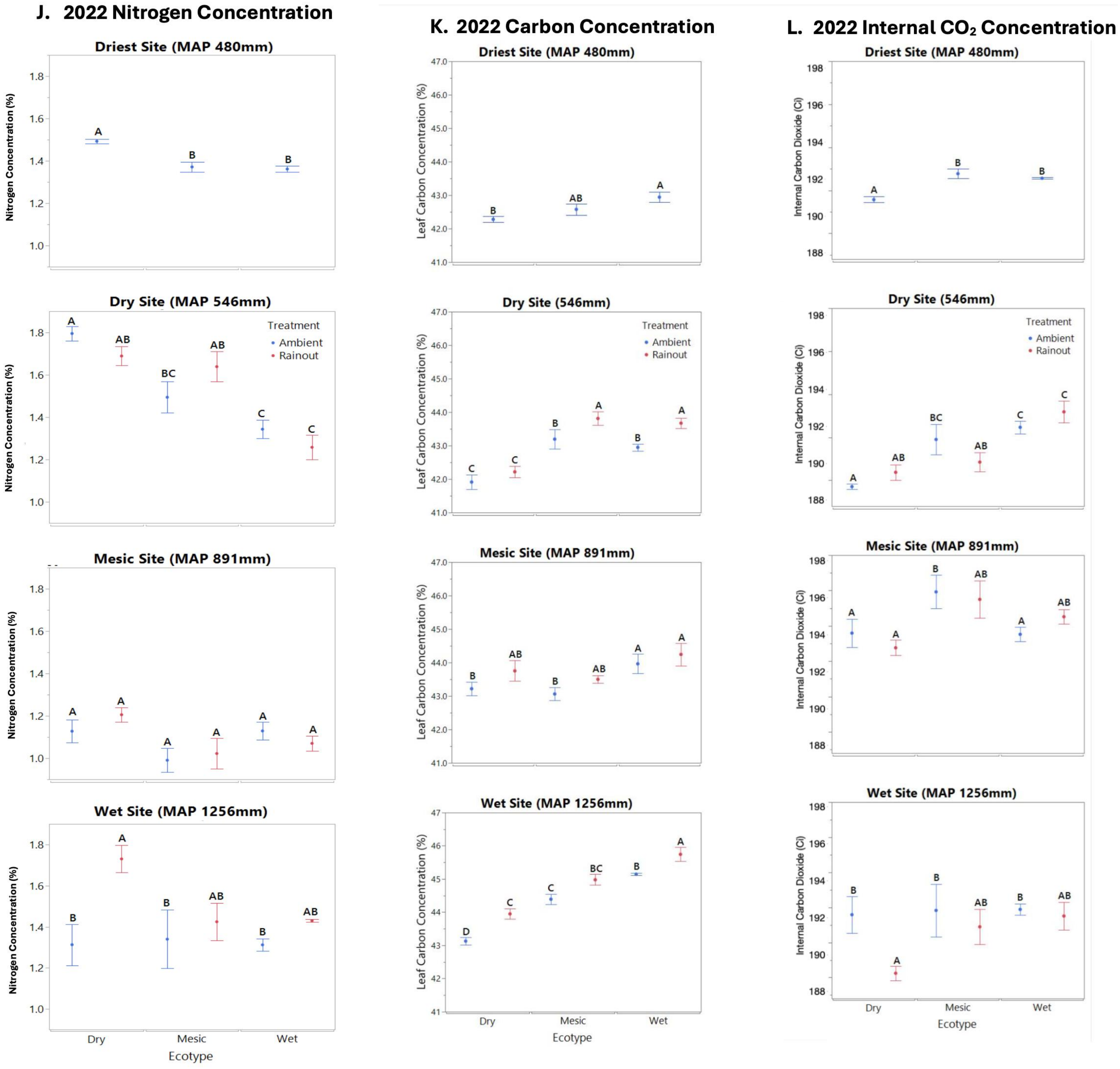
J. 2022 Nitrogen Concentration
K. 2022 Carbon Concentration
L. 2022 Internal CO₂ Concentration
Driest Site (MAP 480mm)
Dry Site (MAP 546mm)
Dry Site (546mm)
Mesic Site (MAP 891mm)
Wet Site (MAP 1256mm)
Treatment
Ambient
Rainout
Nitrogen Concentration (%)
Leaf Carbon Concentration (%)
Internal Carbon Dioxide (Ci)
Dry
Mesic
Wet
Ecotype

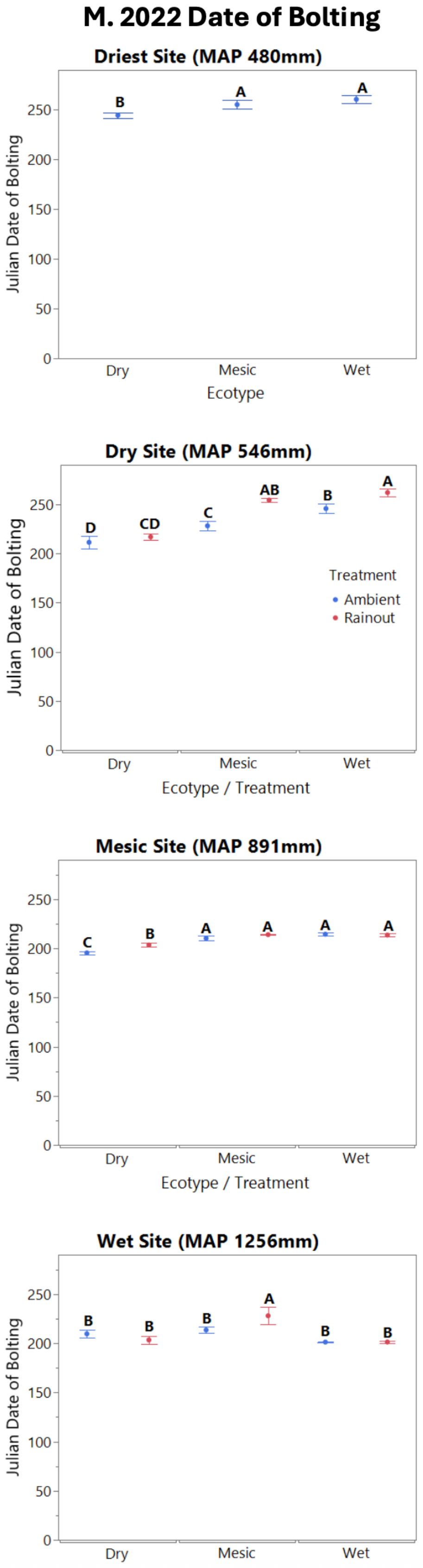

M. 2022 Date of Bolting
Driest Site (MAP 480mm)
Julian Date of Bolting
0
50
100
150
200
250
B
A
A
Dry
Mesic
Wet
Ecotype
Dry Site (MAP 546mm)
Julian Date of Bolting
D
CD
C
AB
B
A
Treatment
Ambient
Rainout
Dry
Mesic
Wet
Ecotype / Treatment
Mesic Site (MAP 891mm)
Julian Date of Bolting
C
B
A
A
A
A
Dry
Mesic
Wet
Ecotype / Treatment
Wet Site (MAP 1256mm)
Julian Date of Bolting
B
B
B
A
B
B
Dry
Mesic
Wet

Supplemental Figure 4. Association between PCA axis 1 and annual precipitation across A. dry, B. mesic and C. wet ecotypes. This shows the relationship between the first principal component (PCA axis 1) and mean annual precipitation (MAP) across sites. Each panel presents a regression analysis to assess the correlation between PCA axis 1 and MAP within each ecotype. Data points represent the mean PCA axis 1 scores for each ecotype, and the regression lines indicate the trend in the relationship between PCA axis 1 and MAP.

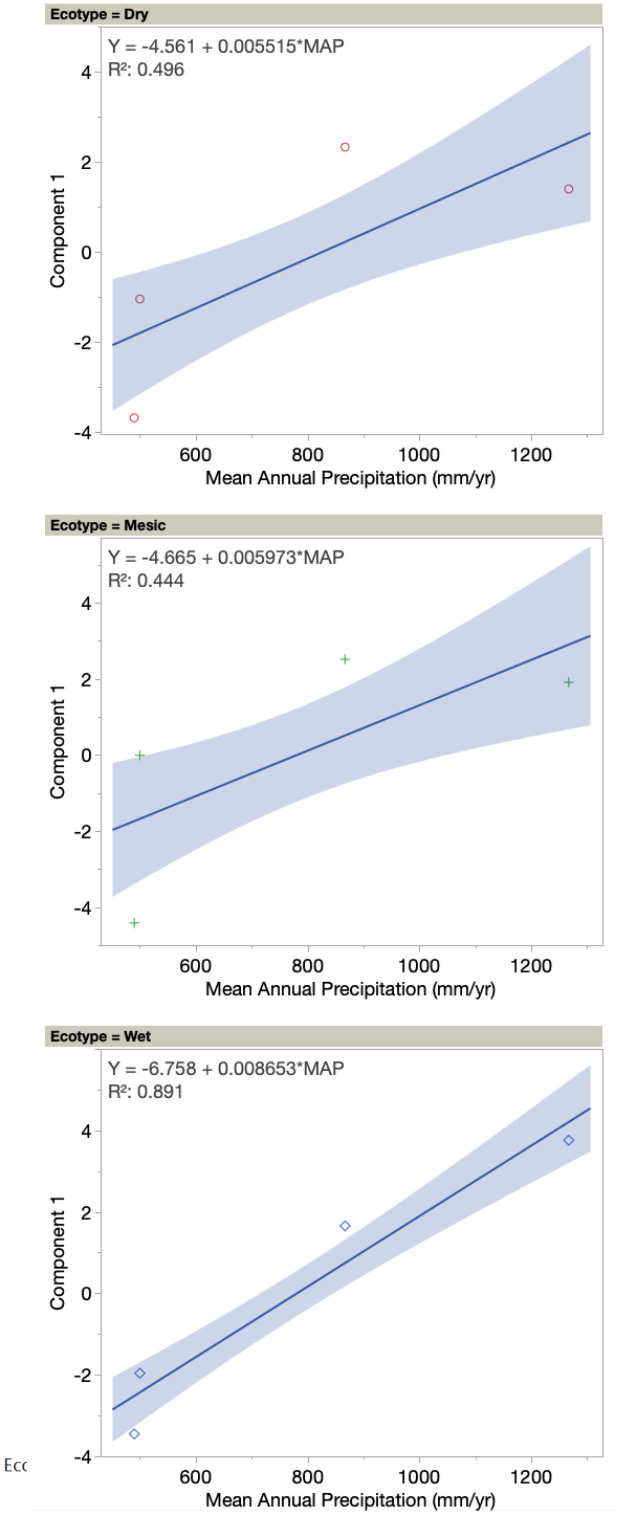

3. Table S1. A. Seed source of the dry, mesic, and wet ecotypes of *A. gerardi* sown into the reciprocal gardens.

| Ecotype | Prairie | Size (ha) | County, State (USA) | Elevation (m) | Soil Type | Latitude | Longitude |
|---|---|---|---|---|---|---|---|
| Dry | Webster Reservoir | 356 | Rooks, KS | 606 | Wakeeney-Hamey Silt Loam | 39.41 | 99.50 |
| Dry | Saline Experimental Range | 880 | Ellis, KS | 641 | Bogue-Armo Clay Loam | 38.76 | 99.14 |
| Dry | Cedar Bluffs Reservoir | 850 | Trego, KS | 688 | Armo Clay Loam | 38.76 | 99.83 |
| Dry | Relict Prairie | 14 | Ellis, KS | 640 | Armo Gravelly Loam | 38.85 | 99.37 |
| Mesic | Konza Prairie | 1557 | Riley, KS | 366 | Benfield Silty Clay Loam | 39.08 | 96.56 |
| Mesic | Tallgrass National Preserve | 4409 | Chase, KS | 392 | Irwin Silty Clay Loam | 38.25 | 96.60 |
| Mesic | Carnahan Cove | 99 | Pottawatomie, KS | 389 | Benfield Silty Clay Loam | 39.34 | 96.62 |
| Mesic | Top of the World Park | 61 | Riley, KS | 379 | Orthents Silty Loam | 39.22 | 96.62 |
| Wet | Desoto Prairie | 0.4 | Jackson, IL | 119 | Orthents Silty Loam | 37.85 | 89.14 |
| Wet | Twelve Mile Prairie | 28 | Effingham, IL | 160 | Cisne Silty Loam | 38.78 | 88.83 |
| Wet | Walters Prairie | 5 | Jasper, IL | 150 | Atlas Silty Clay Loam | 38.92 | 88.19 |
| Wet | Fults Prairie | 214 | Monroe, IL | 215 | Menfro Wetty Clay Loam | 38.17 | 90.19 |

Table S1**.** B. Site information of the reciprocal gardens.

| Reciprocal Garden Site | Elevation (m) | Latitude | Longitude | Mean Annual Temp (C) | Soil Type | Mean Annual Precip (mm) | Annual Precip 2021 (mm) | Annual Precip 2022 (mm) |
|---|---|---|---|---|---|---|---|---|
| Colby, KS, KSU Ag Experimental Station (Thomas Co) | 972 | 39.39 | 101.06 | 10.9 | Ulysses Silt Loam | 480 | 491.1 | 459.8 |
| Hays, KS, KSU Ag Experimental Station (Ellis Co) | 603 | 38.85 | 99.34 | 12.3 | McCook Silt Loam | 546 | 550.8 | 601.1 |
| Manhattan, KS, USDA Plant Material Center (Riley Co) | 315 | 39.19 | 96.58 | 12.8 | Belvue Silt Loam | 891 | 882.9 | 840.0 |
| Carbondale, IL, SIU Ag Research Station (Jackson Co) | 127 | 37.73 | 89.17 | 13.5 | Story Silt Loam | 1256 | 1279.8 | 1196.2 |

**Table S2.** A. Statistical response with main and interactive effect in early season 2022 at p<0.05.

| Response variable | Site | Ecotype | Treatment | Significant Interactions |
|---|---|---|---|---|
| ***Morphological*** | | | | |
| Plant Height | <.0001 | 0.006 | <0.0001 | Site x Eco: <0.0001<br>Site x Trt: 0.0009 |
| Canopy Diameter | <0.0001 | 0.009 | <0.0001 | Eco x Site: 0.01<br>Site x Trt: <0.0001 |
| Leaf Width | <0.0001 | 0.63 | <0.0001 | Eco x Site: <0.0001<br>Eco x Trt x Site: 0.02 |
| Leaf Thickness | <0.0001 | 0.003 | 0.07 | Eco x Site: 0.01<br>Eco x Site x Trt: <0.0001 |
| ***Physiological*** | | | | |
| SPAD | <0.0001 | 0.25 | <0.0001 | Eco x Site: 0.02<br>Trt x Site: <0.0001<br>Trt x Eco x Site: 0.002 |
| Photosynthetic Rate | <0.0001 | <0.0001 | <0.0001 | Site x Eco: 0.014<br>Trt x Site: <0.0001 |
| Transpiration Rate | <0.0001 | 0.01 | 0.16 | Site x Trt: <0.0001<br>Site x Eco x Trt: <0.0001 |
| Stomatal Conductance | <0.0001 | 0.03 | <0.0001 | Trt x Site: <0.0001<br>Eco x Site x Trt: <0.0001 |
| Water Use Efficiency | 0.06 | <0.0001 | 0.03 | Eco x Site: <0.0001<br>Trt x Site: <0.0001<br>Eco x Site x Trt: <0.0001 |
| Internal Carbon Dioxide Concentration | 0.23 | 0.10 | 0.02 | Trt x Site: <0.0001<br>Eco x Site x Trt: <0.0001 |
| ***Reproductive*** | | | | |
| Flowering | <0.0001 | 0.47 | <0.0001 | Eco x Site: <0.0001 |
| Bolting | 0.003 | 0.11 | 0.002 | Eco x Site: 0.04 |

Table S2. B. Statistical response with main and interactive effect in peak season 2022 at p<0.05.

| Response variable | Site | Ecotype | Treatment | Season | Significant Interactions |
|---|---|---|---|---|---|
| ***Morphological*** | | | | | |
| Plant Height | <.0001 | <.0001 | 0.26 | <0.0001 | Site x Eco: <0.0001 |
| Canopy Diameter | <0.0001 | <0.0001 | 0.005 | 0.05 | Eco x Site: 0.0005<br>Eco x Trt x Site :0.03 |
| Leaf Width | 0.02 | <0.0001 | 0.39 | 0.26 | Eco x Site: <0.0001<br>Eco x Trt x Site: 0.007 |
| Nitrogen Concentration | <0.0001 | <0.0001 | 0.06 | N/A | Site x Trt: 0.008<br>Eco x Site: 0.002 |
| Carbon Concentration | <0.0001 | <0.0001 | 0.15 | N/A | Eco x Site: <0.0001 |
| Leaf Thickness | <0.0001 | <0.001 | 0.74 | 0.54 | Eco x Site: <0.0001<br>Trt x Site: 0.02<br>Trt x Site x Eco: 0.02 |
| ***Physiological*** | | | | | |
| SPAD | 0.0001 | <0.0001 | 0.09 | <0.0001 | Seas x Site: <0.0001<br>Eco x Site: <0.0001<br>Trt x Site: 0.008 |
| Photosynthetic Rate | <0.0001 | <0.0001 | 0.0006 | <0.0001 | Site x Eco: 0.01<br>Trt x Site: 0.0008<br>Site x Eco x Trt: 0.01 |
| Stomatal Conductance | 0.01 | <0.0001 | 0.09 | <0.0001 | Seas x Site: 0.001<br>Site x Eco x Trt: 0.01 |
| Transpiration Rate | 0.003 | 0.01 | 0.08 | <0.0001 | Eco x Site: 0.0005<br>Eco x Trt x Site :0.03 |
| Water Use Efficiency | 0.0001 | <0.0001 | 0.002 | <0.0001 | Eco x Site: <0.0001<br>Eco x Trt x Site: 0.007 |

| | | | | | |
|---|---|---|---|---|---|
| Internal Carbon Dioxide Concentration | 0.02 | <0.0001 | 0.06 | 0.04 | Seas x Site: <0.0001<br>Site x Eco x Trt: 0.01 |
| ***Reproductive*** | | | | | |
| Flowering | <0.0001 | <0.0001 | 0.06 | N/A | Eco x Site:0.0005 |
| Bolting | 0.04 | 0.005 | 0.01 | N/A | Eco x Site: 0.01 |
| 2021 Cover | 0.0001 | 0.07 | <0.0001 | N/A | Eco x Site: <0.0001<br>Site x Eco x Trt: 0.03 |
| 2022 Cover | 0.02 | 0.12 | <0.0001 | N/A | Eco x Site: <0.0001<br>Site x Eco x Trt: 0.02 |
| 2021 Biomass | 0.0008 | 0.37 | 0.04 | N/A | Eco x Site: <0.0001<br>Site x Trt:0.02<br>Site x Eco x Trt: 0.01 |
| 2022 Biomass | <0.0001 | 0.10 | 0.07 | N/A | Site x Eco x Trt: 0.02<br>Site x Eco: <0.0001 |

Table S2. C. Statistical response with main and interactive effect in late season 2022 at $p<0.05$.

| **Response variable** | **Site** | **Ecotype** | **Treatment** | **Significant Interactions** |
|---|---|---|---|---|
| ***Morphological*** | | | | |
| Plant Height | <.0001 | 0.02 | 0.29 | Site x Eco: <0.0001<br>Site x Trt: 0.0009<br>Trt x Eco: 0.0009 |
| Canopy Diameter | <0.0001 | <0.0001 | 0.02 | Eco x Site: <0.0001<br>Site x Trt: <0.0001<br>Eco x Trt x Site :0.001 |
| Leaf Width | 0.005 | 0.48 | 0.67 | Eco x Site: <0.0001<br>Eco x Trt x Site: <0.0001 |
| Leaf Thickness | 0.25 | 0.008 | 0.007 | Eco x Site: <0.0001<br>Eco x Trt x Site: <0.0001 |
| Total Biomass | 0.0003 | 0.04 | 0.08 | Eco x Site: <0.0001<br>Eco x Trt x Site: <0.0001 |
| Seed Biomass | 0.001 | 0.24 | 0.08 | Eco x Site: <0.0001<br>Site x Trt: 0.003<br>Eco x Trt x Site: <0.0001 |
| ***Physiological*** | | | | |
| SPAD | 0.04 | 0.017 | <0.0001 | Seas x Site: <0.0001<br>Eco x Site: <0.0001<br>Trt x Site: <0.0001<br>Trt x Eco x Site: 0.014 |
| Photosynthetic Rate | <0.0001 | 0.01 | 0.02 | Site x Eco: <0.0001<br>Trt x Site: <0.0001<br>Site x Eco x Trt: 0.03 |
| Transpiration Rate | <0.0001 | 0.01 | 0.02 | Site x Trt: <0.0001<br>Site x Eco x Trt: <0.0001 |
| Stomatal Conductance | 0.0003 | 0.0002 | <0.0001 | Eco x Site: <0.0001<br>Trt x Site: 0.03<br>Eco x Site x Trt: <0.0001 |
| Water Use Efficiency | 0.006 | 0.04 | 0.15 | Site x Eco: <0.0001<br>Trt x Site: <0.0001 |
| Internal Carbon Dioxide Concentration | 0.45 | 0.15 | 0.05 | Site x Eco: 0.01<br>Trt x Site: 0.002<br>Site x Eco x Trt: <0.0001 |
| ***Reproductive*** | | | | |
| Flowering | <0.0001 | 0.009 | 0.10 | Eco x Site: <0.0001<br>Eco x Site x Trt: 0.014 |
| Bolting | 0.03 | 0.16 | 0.41 | Eco x Site: 0.01<br>Eco x Site x Trt: 0.04 |

**Table S3**. The total number of up- and down-regulated differentially expressed genes at all comparison points and at the individual comparison points for each ecotype (wet or dry), site (wet, dry, or driest), homesite (wet or dry homesite), and treatment (rainout or ambient).

| **Comparison** | **Upregulated genes (padj<0.05)** | **Downregulated genes (padj<0.05)** | **Total differentially expressed genes** |
|---|---|---|---|
| Ecotype (dry vs wet) | 6099 in dry ecotype | 5,179 | 11,278 |
| Dry Homesite | 8404 in dry ecotype | 8820 | 17,224 |
| Wet Homesite | 7,328 in dry ecotype | 8,951 | 16,279 |
| Site, Driest and Dry | 672 in driest site | 683 | 1,355 |
| Site, Dry and Wet | 12,956 in dry site | 11,971 | 24,927 |
| Site, Driest and Wet | 4,631 in driest site | 5,384 | 10,015 |
| Treatment (ambient or rainout) | 221 under rainouts | 187 | 408 |

**Table S4.** Significantly enriched Gene Ontology (GO) terms among differentially expressed genes associated with the main effects of ecotype, site, and rainout treatment. Only GO terms with a false discovery rate (FDR) < 0.05 are included. GO term annotations were obtained using Blast2GO. "u" indicates GO terms enriched among upregulated genes, and "d" indicates enrichment among downregulated genes for each main effect.

| Effect | Gene Function | GO Term |
|---|---|---|
| **Wet Ecotype** | flower bud (u) | GO:0060572 |
| | auxin metabolism processes (u) | GO:0042446 |
| | ATP generation/binding (u) | GO:0005524 |
| | shoot system morphogenesis (u) | GO:0010016 |
| | response to gibberellin (u) | GO:0009739 |
| | gibberellic acid mediated signaling pathway (u) | GO:0009740 |
| | vascular tissue generation (u) | GO:0001944 |
| | RNA modification (u) | GO:0009451 |
| | response to blue light (u) | GO:0009637 |
| | trichome differentiation (u) | GO:0010026 |
| **Dry Ecotype** | root differentiation (u) | GO:0015994 |
| | plant organ senescence (u) | GO:0090693 |
| | regulation of stomatal closure (u) | GO:0015994 |
| | root hair cell differentiation (u) | GO:0048765 |
| | electron transport in photosynthesis (u) | GO:0009767 |
| | guard cell development (u) | GO:0010441 |
| | root epidermis (u) | GO:0010054 |
| | chlorophyll metabolic process (u) | GO:0015994 |
| | stomatal complex development (u) | GO:0010374 |
| | primary root growth (u) | GO:0080022 |
| | defense response (u) | GO:0006952 |
| | regulation of photoperiodism (u) | GO:0048573 |
| | flowering (u) | GO:0048573 |
| **Wet Site** | biosynthetic process (u) | GO:0008152 |
| | metabolic processes (u) | GO:0008152 |
| | regulation of biosynthetic process (u) | GO:0009889 |
| | salicylic acid mediated signaling pathway (u) | GO:2000031 |
| | cellular response to amino acid starvation (u) | GO:0034198 |
| | abscisic acid catabolic process (d) | GO:0046345 |
| | auxin response (u) | GO:0009733 |
| | shoot system meristem (u) | GO:0010016 |
| | reproductive bud (u) | GO:0009908 |
| | vascular system (u) | GO:0010051 |
| | chloroplast organization (u) | GO:0015994 |
| **Dry Site** | primary root growth (u) | GO:0010102 |
| | regulation of metabolism (d) | GO:0019222 |
| | electron transporter (u) | GO:0009772 |
| | defense responses (u) | GO:0006952 |
| | DNA-binding transcription factor activity (d) | GO:0003700 |
| | regulation of transcription (u) | GO:0006357 |
| | stress response (u) | GO:0006952 |

| | | |
|---|---|---|
| | DNA repair (u)<br>root cap of primary root (u)<br>ATP binding (u)<br>hydrolase activity (u)<br>lateral root growth (u) | GO:0006281<br>GO:0048829<br>GO:0005524<br>GO:0016787<br>GO:0010102 |
| **Driest Site** | Defense responses (u)<br>DNA-binding transcription factor activity (d)<br>cellular copper ion homeostasis (u)<br>abscisic acid transport (u)<br>programmed cell death (u)<br>lateral root tip (u)<br>chitinase activity (u)<br>flower bud (d) | GO:0006952<br>GO:0003700<br>GO:0006878<br>GO:0080168<br>GO:0012501<br>GO:0048527<br>GO:0004568<br>GO:0048573 |
| **Wet Homesite** | apical meristem growth (u)<br>seed maturation (u)<br>flower growth (u)<br>hormone metabolism processes (u)<br>shoot system morphogenesis (u)<br>response to gibberellin (u)<br>hormone extracellular transport (u)<br>gibberellic acid mediated signaling pathway (u)<br>response to hypoxia (u) | GO:0040008<br>GO:0048573<br>GO:0048573<br>GO:0006106<br>GO:0010016<br>GO:0009739<br>GO:0006858<br>GO:0009740<br>GO:0001666 |
| **Dry Homesite** | pollen development (u)<br>regulation of metabolism (u)<br>electron transporter (u)<br>transferring electrons for photosynthesis activity (u)<br>guard cell development (u)<br>photosynthesis (u)<br>root epidermis (u)<br>abscisic acid-activated signaling pathway (u)<br>chloroplast modification (u) | GO:0010431<br>GO:0019222<br>GO:0009772<br>GO:0009767<br>GO:0010377<br>GO:0009658<br>GO:0010053<br>GO:0080168<br>GO:0009658 |
| **Rainout** | root apical meristem (u)<br>chloroplast modification (u)<br>DNA damage checkpoint signaling (u)<br>heat shock protein signaling (u)<br>cellular water homeostasis (u)<br>negative regulation of meristematic growth (u)<br>root apical meristem (u)<br>acyl-CoA dehydrogenase activity (u)<br>lateral root cap (u) | GO:0048829<br>GO:0009507<br>GO:0000077<br>GO:0031072<br>GO:0009992<br>GO:0010075<br>GO:0010071<br>GO:0003995<br>GO:0048829 |
| **Ambient** | vascular growth (u)<br>first order inflorescence axis (u)<br>stem epidermis (u)<br>hormone transport (u)<br>regulation of metabolism (u)<br>positive regulation of seed maturation (u)<br>intracellular monoatomic ion homeostasis (u)<br>regulation of growth (u) | GO:0001944<br>GO:0009672<br>GO:0090558<br>GO:0009914<br>GO:0019222<br>GO:2000693<br>GO:0006873<br>GO:0040008 |

**Table S5.** List of selected candidate genes identified across ecotypes, sites, treatment, and homesite. The table includes gene annotations, associated Gene Ontology (GO) terms, and fold change values indicating the direction and magnitude of differential expression.

| Effect | Gene | Annotation | GO Term | Fold Change |
|---|---|---|---|---|
| **Ecotype** | 06BG063200 | abscisic stress-ripening | 0006950 | +3.57 |
| Dry Ecotype | 05BG130400 | 3-ketoacyl-CoA synthase | 0006633 | +5.47 |
| | 04BG198300 | *ARF16* | 0005524 | +6.29 |
| | 01CG411500 | chloroplastic drought-induced stress protein | 0045454 | +6.34 |
| | 02CG047700 | Chlorophyll A-B binding family protein | 0005515 | +6.11 |
| | 09CG201100 | heat shock protein 17.4 | 0006950 | +6.92 |
| | 04AG294100 | chloroplasts 55-II | 0016491 | +5.44 |
| | 04BG127400 | salt stress root protein RS1 | 0051716 | +4.02 |
| | 07BG164800 | rotamase FKBP 1 | 0006457 | +6.47 |
| | 05AG094300 | Heat shock protein 22H | 0005515 | +5.06 |
| | 01CG303500 | SAUR-like auxin-responsive protein | 0009733 | +4.44 |
| | 09BG190100 | indole-3-butyric acid response 1 | 0008667 | +4.39 |
| | 10CG027000 | early nodulin 93 | 0016757 | +4.31 |
| | 10AG161400 | multiprotein bridging factor 1C | 0043565 | +4.24 |
| **Ecotype** | 02AG195700 | gibberellin receptor GID1L2 | 0016491 | +4.52 |
| Wet Ecotype | 03CG144300 | auxin response factor 75 | 0006281 | +3.15 |
| | 05BG011300 | WRKY64 | 0006355 | +4.11 |
| | 03BG273300 | gibberellin 2-oxidase 1 | 0005506 | +3.90 |
| | 01CG090600 | BEL1-like homeodomain 4 | 0006355 | +3.23 |
| | 01CG314300 | myb domain protein 94 | 0003677 | +4.10 |
| | 08CG126900 | Auxin-responsive Aux/IAA gene family | 0005634 | +4.46 |
| | 03CG275000 | HVT1 Vascular Tissue Tapetum | 0005524 | +4.09 |
| | 10BG199700 | auxin response factor 16 | 0005634 | +3.20 |
| | 01CG072700 | GASR3 | 0008080 | +4.78 |
| | 02AG291100 | gibberellin-responsive protein | 0008080 | +3.06 |
| **Site** | 03AG057500 | FASCICLIN-like arabinogalactan-protein 11 | 0006351 | -3.11 |
| Dry Site | 01AG088000 | senescence-associated gene 12 | 0008234 | +4.20 |
| | 04BG038400 | STRUBBELIG-RECEPTOR FAMILY 1 | 0008324 | +3.87 |
| | 01AG264300 | transcription factor MYC7E | 0004559 | +3.01 |
| | 08BG013400 | GRAS transcription factor | 0055114 | -4.15 |
| | 06BG118800 | TRS120 | 0003676 | -3.64 |
| | 07CG097100 | phototropin 2 | 0004672 | +3.05 |
| | 05CG174200 | Stigma-specific Stig1 | 0003723 | -4.09 |
| | 02CG360600 | Ethylene insensitive 3 | 0003700 | +4.60 |
| | 03BG256400 | senescence-associated protein | 0050662 | +3.22 |
| | 04AG218300 | cold-regulated 47 | 0009415 | -4.06 |
| | 10AG102800 | heat shock protein DnaJ | 0008168 | +3.44 |
| | 06AG010600 | heat shock protein 90.1 | 0006457 | +6.13 |
| **Site** | 10AG189900 | alpha/beta-Hydrolases | 0008236 | -9.06 |
| Wet Site | 02AG172700 | auxin-repressed protein | 0009733 | -6.14 |
| | 02AG115500 | aquaporin protein 2 | 0005215 | +3.25 |
| | 03CG094900 | DNAJ Heat Shock 20 | 0005743 | +3.12 |
| | 09AG005900 | auxin-induced protein 5NG4 | 0016021 | +5.14 |
| | 07BG171400 | ascorbate peroxidase 6 | 0055114 | -3.21 |
| | 05BG105600 | BRASSINOSTEROID INSENSITIVE 1 | 0005515 | +3.81 |
| | 10AG218700 | Amidase 1 | 0016884 | +3.31 |
| | 02BG022000 | photosynthetic electron transfer A | 0009055 | +5.44 |
| | 01AG387800 | MYB family transcription factor | 0003677 | +3.74 |
| | 04AG206000 | aldehyde dehydrogenase 3F1 | 0004030 | -3.11 |
| **Site** | 03BG150100 | hypoxia-responsive family protein | 0020037 | -6.94 |
| Driest Site | 07CG071600 | photosystem II subunit | 0009523 | +4.78 |
| | 04AG268100 | Chloroplast ATP synthase delta chain | 0046933 | -3.75 |
| | 04BG108300 | Flowering Locus T gene | 0006468 | -4.64 |
| | 10AG184500 | cytochrome P450 | 0005506 | +3.37 |
| | 06CG013300 | heat shock protein 90.1 | 0006457 | -4.72 |

| | | | | |
|---|---|---|---|---|
| | 04AG254400 | ATP-BINDING TRANSPORTER ABC | 0016020 | +3.45 |
| | 02AG115600 | aquaporin protein | 0006810 | +6.73 |
| | 02AG053400 | SHOOT1 protein | 0005515 | +6.09 |
| | 03BG135200 | universal stress protein domain | 0006950 | +5.18 |
| | 05BG075500 | gibberellin receptor G3 | 0055114 | -4.49 |
| **Rainout** Main effect | 05AG211000 | stachyose synthase | 0003824 | +5.28 |
| | 02CG154300 | armadillo/beta-catenin | 0005488 | +5.87 |
| | 03BG099300 | stress responsive protein | 0006355 | +6.27 |
| | 03BG099200 | no apical meristem protein | 0003677 | +3.40 |
| | 01AG088000 | senescence-associated gene 12 | 0008234 | +3.44 |
| | 04CG017700 | DROUGHT SENSITIVE 1 | 0005506 | +3.13 |
| | 07AG141100 | panicle inflorescence | 0005515 | +3.79 |
| | 03BG198000 | stress responsive A/B Barrel | 0005515 | +3.75 |
| | 10BG085100 | annexin 7 | 0005509 | +4.03 |
| **Ambient** Main effect | 03BG392900 | glutathione S-transferase TAU 8 | 0005515 | -5.52 |
| | 03CG305900 | NAC Transcription Factor | 0006355 | +5.61 |
| | 04CG017700 | Abscisic acid-responsive (TB2/DP1, HVA22) | 0005515 | -4.98 |
| | 03CG188600 | MADS14 | 0003676 | -3.05 |
| | 03BG161400 | EX07A2 | 0000145 | -3.14 |
| | 03CG380400 | ALOG2 Shoot Apical Meristem | 0003676 | +3.40 |
| | 03CG182000 | FLOWERING LOCUS C | 0003676 | +3.45 |
| | 03BG381500 | Haloacid dehalogenase (HAD) | 0000287 | -3.51 |
| | 03CG182300 | Albino Leaf 2 | 0003676 | -3.15 |
| | 09CG098800 | Dehydrin | 0009415 | -3.91 |
| **Homesite** Dry Ecotype x Dry Site | 03AG271600 | Flowering Locus T | 0003676 | +5.22 |
| | 08BG118900 | FRIGIDA-like protein | 0005509 | +3.17 |
| | 06CG133000 | Photosystem II reaction center PsbP protein | 0005488 | +3.26 |
| | 01BG420000 | heat shock protein 17.4 | 0005515 | +4.47 |
| | 10BG141200 | DAWDLE | 0005515 | +6.52 |
| | 04CG269300 | heat shock 22 kDa protein | 0005515 | +5.86 |
| | 04AG159700 | STRUBBELIG-RECEPTOR FAMILY 7 | 0004713 | +6.26 |
| | 02BG224700 | Chaperonin | 0006457 | +6.10 |
| | 02BG224200 | photosystem I light harvesting complex gene 6 | 0016020 | +6.46 |
| | 05AG023100 | acyl-CoA synthetase protein | 0008152 | +5.27 |
| | 06AG091500 | auxin response factor 1 | 0005634 | +3.15 |
| | 07BG137500 | EXPANSIN-A1 | 0009664 | +4.76 |
| | 03BG188700 | Chaperone DnaJ | 0005515 | +4.69 |
| | 02AG130800 | flowering time control protein FCA | 0003676 | +3.12 |
| **Homesite** Wet Ecotype x Wet Site | 09AG131900 | gibberellin receptor GID1L2 | 0016787 | +4.49 |
| | 02CG241200 | SAUR-like auxin-responsive protein | 0009733 | +3.71 |
| | 05BG170800 | SCARECROW | 0005202 | +3.43 |
| | 01AG142600 | Auxin-responsive gene family | 0009733 | +4.11 |
| | 08BG148200 | senescence-related gene 1 | 0055114 | -3.55 |
| | 03CG003900 | growth-regulating factor 5 | 0005524 | -3.58 |
| | 03CG122300 | Plant regulator RWP-RK family protein | 0003677 | +3.99 |
| | 03CG017400 | HMG box protein with ARID/BRIGHT | 0003676 | +4.12 |
| | 03CG014300 | histone H3 | 0003676 | -3.45 |
| | 06AG110600 | GASR4 - Gibberellin-regulated GASA | 0008080 | +3.07 |

**Table S6.** The DEGs co-expressed with plant phenotype and their candidate genes associated, annotation, WGCNA cluster number, and GO term. Phenotype abbreviations: Ph= Photosynthetic Rate, Co= Stomatal conductance, Tr=Transpiration Rate, Wu= Water Use Efficiency, Sp= SPAD, Ci= Internal Carbon Dioxide Concentration, Bw=Blade Width, Rg= Relative Growth Rate, He=Height, Th= Leaf Thickness, Di= Canopy Diameter, Sb=Seed biomass, Tb=Total Biomass, Nc= Leaf Nitrogen Concentration, Cc= Leaf Carbon Concentration, Bo=Bolting, and Fl=Flowering.

| Effect | Gene | Annotation | Gene Ontology | Phenotype(s) code | Cluster (Fig S10) | Fold Change |
|---|---|---|---|---|---|---|
| **Ecotype** | 04BG198300 | *ARF16* | 0005524 | Tr, Ci, He, Rg, Cc | 33 | +6.29 |
| Dry | 09CG180800 | chlorophyll A-B binding protein | 0009765 | Tr, Ci, He | 26 | +4.40 |
| Ecotype | 02CG351100 | drought induced protein 19 | 0003676 | Ci, Wi, Nc, Cc | 26 | +3.98 |
| | 01CG411500 | chloroplastic drought-induced stress protein | 0045454 | Tr, Ci, He | 26 | +3.34 |
| | 03AG366700 | thylakoid lumenal protein, chloroplast precursor | 0008152 | Tr, Ci, He, Cc | 33 | +3.17 |
| | 01CG308200 | HVA22 | 0045454 | Tr, Ci, He, Cc | 33 | +3.11 |
| | 08AG143200 | SNF2 | 0005524 | Wi, Th, Tb | 12 | -2.99 |
| | 08AG043900 | LPTL | 0008270 | Ci, Wi, Nc, Cc | 19 | +3.36 |
| | 08AG131200 | Calmodulin | 0005516 | Wi | 1 | +3.47 |
| | 04AG325000 | Bzip txn factor | 0043565 | Tr, Ci, He | 26 | +3.55 |
| | 09AG159900 | Thioredoxin | 0045454 | Tr, Ci, He, Cc | 33 | +3.87 |
| | 08AG074800 | DREB2A | 0046872 | Ci, He | 13 | +4.15 |
| | 08AG174800 | Chaperonin 60 | 0042026 | Tr, Ci, He, Rg, Cc | 33 | -3.66 |
| **Ecotype** | 10BG190500 | PAPA-1-like conserved family protein | 0031011 | Ph, Tr, Wu, Rg, He, Di, Sb, Tb, Cc, Bo, Fl | 8 | +3.61 |
| Wet | 01AG027500 | GASR3 - Gibberellin-regulated family protein | 0016491 | Ph, Tr, Wu, Rg, He, Di, Sb, Tb, Cc | 29 | +3.98 |
| Ecotype | 03AG306700 | cellular response to gibberellin | 0016491 | Ph, Tr, Wu, Rg, He, Di, Sb, Tb, Cc | 29 | +3.37 |
| | 03BG134600 | SH2 motif | 0006139 | Ph, Tr, Wu, Rg, He, Di, Sb, Tb, Cc, Bo, Fl | 8 | +3.67 |
| | 03CG144300 | ARF-75 | 0006281 | Ph, Co Tr, Wu, Rg, Sb, Tb, Cc, Bo, Fl | 11 | +3.15 |
| | 10AG219900 | gibberellin 2-oxidase 1 | 0030896 | Ph, Co Tr, Wu, Rg, Sb, Tb, Cc, Bo, Fl | 30 | +4.11 |
| | 04AG308700 | basic leucine zipper 9 Transcription Factor | 0003700 | Di, Cc, Nc | 30 | +3.45 |
| | 10BG070600 | nudix hydrolase homolog 3 | 0016787 | Co, Tr, Wu, Ci, Rg, Sb, Tb, Bo, Fl | 3 | +3.97 |
| | 10BG153500 | 14-3-3 Protein | 0019904 | Ph, Co Tr, Wu, Rg, Sb, Tb, Cc, Bo, Fl | 9 | -3.59 |
| | 10BG150800 | OMP85 family protein | 0019867 | Ph, Co Tr, Wu, Rg, Sb, Tb, Cc, Bo, Fl | 11 | +5.37 |
| | 03CG144300 | PHABULOSA | 0032784 | Ph, Co Tr, Wu, Rg, Sb, Tb, Cc, Bo, Fl | 11 | +3.88 |
| | 10BG155600 | B4FSM0, **Anther-specific protein APG** | 0048653 | Ph, Co Tr, Wu, Rg, Sb, Tb, Cc, Bo, Fl | 11 | +3.06 |
| **Site** | 07AG194100 | DROUGHT SENSITIVE 1 | 0005515 | Ph, Co, Rg | 5 | +3.83 |
| Dry Site | 06CG148400 | DNAJ heat shock N-terminal protein | 0003677 | He, Sb | 11 | +3.45 |
| | 03BG201200 | growth inhibition & differentiation protein 88 | 0005506 | Co, Nc, Cc, Bo | 4 | +4.24 |
| | 04AG059600 | XPB2 | 0003677 | Ph, Co, Rg | 5 | +3.97 |
| | 07AG184300 | heat shock protein STI | 0005515 | Co, Nc, Cc, Bo | 4 | +3.15 |
| | 03CG129700 | Drought-responsive family protein | 0003676 | Co, Sb | 6 | +3.57 |
| | 05CG083100 | CLAVATA1 | 0004672 | Co, Sb | 6 | +4.77 |
| | 05CG144800 | MRH1 | 0004672 | Th | 8 | -3.56 |
| | 05CG160700 | DUF26 | 0004672 | Th | 8 | +3.68 |
| | 05CG091400 | RPS2 protein | 0003684 | He, Sb | 11 | +4.11 |
| | 03CG062100 | heat shock protein ATPase | 0001671 | He, Sb | 11 | +3.07 |
| | 05CG022600 | STRUBBELIG-RECEPTOR FAMILY 7 | 0004672 | Sp | 18 | +3.95 |
| **Site** | 03BG175900 | phytochrome C | 0000155 | Nc, Cc | 2 | +3.12 |
| Wet Site | 08AG163200 | growth-regulating factor | 0005524 | He, Di | 8 | +3.07 |
| | 03BG285900 | flowering time control protein FCA | 0000166 | He, Di | 8 | +3.71 |
| | 04BG018000 | chloroplast splicing factor CRS1 | 0003723 | He, Di | 8 | +3.50 |
| | 04CG101100 | DWARF1 | 0003824 | Nc, Cc | 2 | -3.45 |
| | 04BG203700 | light-mediated development protein DET1 | 0003824 | He, Di | 8 | +4.97 |
| | 08AG138500 | PSI-N | 0005516 | He | 5 | +4.22 |
| | 04CG097900 | DIMINUTO | 0003824 | He, Di | 8 | -3.69 |
| | 03BG173900 | ethylene receptor | 0000155 | He, Di | 8 | +3.99 |
| | 03BG271200 | translation initiation factor 3B1 | 0000166 | He, Di | 8 | +3.95 |
| | 03CG181700 | RNA Binding flowering time control | 0003676 | Bo | 10 | -3.88 |
| | 03BG247900 | RRM/RBD/RNP motifs family protein | 0000166 | Wi | 15 | +3.22 |
| | 03BG361500 | beta-amylase 7 | 0000272 | Bo | 16 | -3.17 |
| **Site** | 04CG002300 | 3-ketoacyl-CoA synthase precursor | 0003824 | Nc, Cc | 5 | +4.94 |
| Driest | 03CG339100 | OsLonP2 protease | 0003684 | He, Di | 12 | +3.66 |
| Site | 03CG336100 | AP2/B3-like transcriptional factor protein | 0003677 | He, Di | 12 | +3.08 |
| | 04CG023900 | hydrolase, alpha/beta fold family | 0003824 | Bo | 14 | +3.96 |
| | 03CG334200 | GROWTH INHIBITION | 0000166 | Nc | 22 | +4.88 |
| | 03CG331900 | chalcone synthase | 0003824 | He, Cc | 24 | -3.15 |
| | 04CG056200 | OPC-8:0 CoA ligase1 | 0003824 | Rg, He, Di, Bo | 26 | +3.88 |
| | 03BG188900 | two-component response regulator | 0000160 | He, Di | 12 | +3.55 |
| | 04CG016400 | methyl-CpG binding domain | 0003677 | He, Di | 12 | +3.96 |

| | | | | | | |
|---|---|---|---|---|---|---|
| **Rainout** | 06BG244400 | aberrant root formation protein 4 | 0005488 | Ph, Co, Tr, Wu, Rg, He, Tb, Nc, Bo | 3 | +3.93 |
| Main | 07AG210300 | stress responsive A/B Barrel | 0005515 | Ph, Co, Tr, Wu, Rg, He, Tb, Nc, Bo | 3 | +3.75 |
| effect | 07AG141100 | DROUGHT SENSITIVE 1 | 0005506 | Ph, Co, Tr, Wu, Rg, He, Tb, Nc, Bo | 3 | +3.13 |
| | 06CG056900 | Drought-responsive family protein | 0005515 | Ph, Co, Tr, Wu, Rg, He, Tb, Nc, Bo | 3 | +3.04 |
| | 06BG244100 | Impaired sucrose induction 1 | 0005488 | Ph, Co, Tr, Wu, Rg, He, Tb, Nc, Bo | 3 | +4.99 |
| | 07AG210300 | stress-inducible protein | 0000166 | Ph, Co, Tr, Wu, Rg, He, Tb, Nc, Bo | 3 | +3.67 |
| | 03BG198000 | panicle inflorescence | 0005515 | Ph, Co, Tr, Wu, Rg, He, Tb, Nc, Bo | 3 | +3.79 |
| | 06BG244300 | ALF4-1 | 0005488 | Ph, Co, Tr, Wu, Rg, He, Tb, Nc, Bo | 3 | -3.97 |
| **Ambient** | 03CG182300 | Albino Leaf 2 | 0003676 | Ph, Tr, Ci, Bo, Fl | 3 | -3.15 |
| Main | 03CG182000 | FLOWERING LOCUS C | 0003676 | Sp | 1 | +3.45 |
| effect | 03BG381500 | Haloacid dehalogenase (HAD) | 0000287 | Sp | 1 | -3.51 |
| | 03BG258200 | SC35 | 0000166 | Sp | 1 | +3.94 |
| | 03CG188600 | MADS14 | 0003676 | Tb | 7 | -3.05 |
| | 03BG161400 | EX07A2 | 0000145 | Sp | 1 | -3.14 |
| | 03CG380400 | ALOG2 Shoot Apical Meristem | 0003676 | Ph, Tr, Ci, Bo, Fl | 3 | +3.40 |
| | 03CG178300 | BFN1 | 0003676 | Ph, Tr, Ci, Bo, Fl | 3 | -3.66 |
| **Homesit** | 08BG113900 | expansin B2 | 0005576 | Tr, Wu, Ci, Rg, Tb, Nc, Cc, Bo, Fl | 10 | +3.08 |
| **e** | 08BG118900 | FRIGIDA-like protein | 0005509 | Wu, Wi, Tb, Nc, Cc, Bo | 16 | +3.17 |
| Dry | 06CG133000 | Photosystem II reaction center PsbP protein | 0005488 | Ph, Co, Wu, Ci, Rg | 9 | +3.26 |
| Ecotype | 06BG242400 | aberrant root formation P4 | 0005525 | St, He, Sb | 17 | +3.02 |
| x Dry | 08BG071800 | Root hair defective 5 | 0005525 | Tr, Wu, Ci, Rg, Tb, Nc, Cc, Bo, Fl | 10 | -3.94 |
| Site | 08BG159100 | auxin response factor 8 | 0005634 | Ph, Co, Wu, Ci, Rg | 9 | +3.91 |
| | 08AG085500 | chloroplast signal recognition (CAO) | 0005515 | Ph, Co, Wu, Ci, Rg | 9 | +4.12 |
| | 08BG154800 | transcription factor jumonji | 0005634 | Sp, Th | 20 | +2.05 |
| | 08AG138400 | chloroplast PSI-N | 0005516 | Sp, Th | 20 | +3.47 |
| | 08BG154700 | chloroplast import apparatus 2 | 0005509 | Ph, Co, Rg | 6 | +3.46 |
| | 08BG138600 | CCT/B-box zinc finger protein | 0005622 | Co, He, Sb | 17 | +3.87 |
| | 08BG130200 | SNARE-like superfamily protein | 0005622 | Ph, Co, Rg | 6 | +3.95 |
| | 08BG113800 | EXPANSIN-A1 | 0005576 | Tr, Wu, Ci, Rg, Tb, Nc, Cc, Bo, Fl | 10 | +3.66 |
| **Homesit** | 03CG374700 | myb family transcription factor | 0003677 | Bo | 2 | +3.01 |
| **e** | 04AG186600 | cup-shaped cotyledon 3 | 0000413 | Ph, Co, Tr, Wu, Ci, Nc, Cc, Bo, Sb, Fl | 23 | +3.42 |
| Wet | 03CG190500 | PINHEAD | 0003676 | Ph, Co, Tr, Wu, Sp, Rg, He, Di, Fl | 25 | +4.05 |
| Ecotype | 06CG068600 | gibberellin 2-beta-dioxygenase 7 | 0003677 | Ph, Co, Tr, Wu, Rg, He, Di, Sb, Tb, Bo, Fl | 6 | +3.96 |
| x Wet | 06CG071100 | gibberellin 20 oxidase 1 | 0003676 | Ph, Co, Tr, Wu, Rg, He, Di, Sb, Tb, Nc, Cc, | 5 | +3.66 |
| Site | 03CG351600 | nitrogen metabolism transcription factor | 0003677 | Nc | 26 | +5.16 |
| | 03CG003900 | growth-regulating factor 5 | 0005524 | Nc | 26 | -3.58 |
| | 03CG122300 | Plant regulator RWP-RK family protein | 0003677 | Nc | 26 | +3.99 |
| | 03CG017400 | HMG box protein with ARID/BRIGHT | 0003676 | Nc | 26 | +4.12 |
| | 03CG014300 | histone H3 | 0003676 | Nc | 26 | -3.45 |
| | 03CG190500 | DCL3 | 0003676 | Nc | 25 | -4.00 |
| | 03CG326800 | TRF-like 6 | 0003677 | Ph, Co, Tr, Wu, Sp, Rg, He, Di, Fl | 23 | +3.22 |
| | 04AG185300 | no apical meristem protein | 0000413 | Ph, Co, Tr, Wu, Ci, Nc, Cc, Bo, Sb, Fl | 23 | -3.76 |
| | 04AG182900 | VASCULAR-RELATED NAC-DOMAIN 6 | 0003676 | Ph, Co, Tr, Wu, Ci, Nc, Cc, Bo, Sb, Fl | 23 | +4.21 |
| | 03CG242200 | cyclophilin 38 | 0000413 | Ph, Co, Tr, Wu, Ci, Nc, Cc, Bo, Sb, Fl | 23 | +3.28 |
| | 03CG402000 | growth-regulating factor 3 | 0005524 | Ph, Co, Tr, Wu, Ci, Nc, Cc, Bo, Sb, Fl | 23 | +3.83 |
| | 03CG127400 | early flowering 6 | 0003676 | Ph, Co, Tr, Wu, Ci, Nc, Cc, Bo, Sb, Fl | 23 | +3.26 |
| | 03CG388900 | ethylene-responsive transcription factor | 0003676 | Di | 17 | +4.79 |

5.

**Methods S1: Extraction and Mapping**

Total RNA was isolated and purified using the QIAGEN RNeasy kit according to the manufacturer's recommended protocol for plants. The concentration and purity of total RNA was evaluated on the Nanodrop Spectrophotometer (Thermo Scientific). The intactness of RNA samples was assessed using the Agilent Bioanalyzer 2100 (Agilent Technologies). To eliminate genomic DNA contamination of the RNA samples were treated with RNase-Free DNase I (Qiagen). cDNA libraries were constructed from messenger RNA and prepared for sequencing on the Illumina NovaSeq 600 in two lanes. The pooled libraries were then loaded onto Illumina NovaSeq 600 at a concentration of 14 pM and sequenced for 100 cycles from each end of the fragments plus 7 cycles for the index read according to the manufacturer's instructions (Illumina, San Diego, CA). Raw files were converted to fastq files with Casava 1.8.2. Average quality scores were ≥30 across sequencing cycles and reads were 150 bp in length.

Prior to mapping, raw reads were processed using Trimmomatic v.0.33 (Bolger et al., 2014) to remove adapters and the quality of resulting trimmed and cleaned reads was assessed using *FastQC v0.11.9 (*Andrews, 2010). Reads were then mapped to the assembly version of the *Andropogon gerardi* Hap2 v1.1 genome (Joint Genomics Institute, 2023) using the splice-aware mapper Hitsat2 v.2.0.0 (Wen, 2017). We used HiSat2 (Wen, 2017) to index the reference genome and each of the individual samples was mapped to the indexed reference genome. Next, gene counts were obtained using the *feature Counts (*Liao et al., 2014) procedure as implemented in HISat2 (Wen, 2017).
